# Solution Phase Protein Adsorption to ss(GT)_15_-DNA Wrapped Single Walled Carbon Nanotubes

**DOI:** 10.64898/2026.05.18.725765

**Authors:** Gabriel Sánchez-Velázquez, Thomas K. Porter, Lucas Ospina, Ali A. Alizadehmojarad, Wonjun Yim, Xiaoyi Wang, Michael S. Strano

**Affiliations:** Department of Chemical Engineering, Massachusetts Institute of Technology, 77 Massachusetts Avenue, Cambridge, Massachusetts 02139, United States

**Keywords:** metalloprotein, single-walled carbon nanotubes, photophysics, protein binding

## Abstract

Proteins in solution adsorb to the corona of nanoparticles such as single-walled carbon nanotubes (SWCNTs), but these interactions are difficult to predict and analyze due to ambiguities in the structure of the latter. In this work, we employ ss(GT)_15_-DNA wrapped SWCNTs, a commonly used fluorescent sensor construct, to examine protein adsorption by quantifying binding dissociation constants and characterizing the corresponding photophysical effects. A library of 20 proteins are used to evaluate adsorption-induced changes in photoluminescence (PL) intensity (ΔI/I_0_) and emission wavelength upon solution phase binding. We find that 15 proteins produce monotonic dose–response behavior well described using a single-site Langmuir model. Alternatively, five proteins exhibited more complex, non-monotonic behavior consistent with a two-step binding model representing protein–protein interactions coupled to adsorption. The study reveals that metalloproteins, which comprised 12 of the 20 proteins in the library, induced greater PL quenching compared with metal-free proteins for this system, with maximum binding-associated quenching (ΔI/I_0_) of 94% for metalloproteins versus 20% for metal-free proteins. For metalloproteins, we introduce a proximity-based quenching framework in which protein size provides a coarse proxy for cofactor–SWCNT separation, offering a mechanistic interpretation of the observed quenching variation across proteins. Together, these results establish the use of metal coordination sites, such as those in metalloproteins, to assist the transduction of certain nanoparticle fluorescent sensors, helping with sensor probe design and interpretation in biological environments.

## Introduction

Upon introduction into complex biological media, proteins rapidly adsorb to nanoparticles forming the basis for drug delivery, biocatalytic and biosensor technologies.^1,2^ The nanoparticle surface is typically decorated with specific chemistry to assist particle dispersion and protein interactions in a layer described as the nanoparticle corona. On adsorption, proteins can add to this corona forming a complex driven by electrostatic, hydrophobic, hydrogen-bonding, and π–π interactions. The process is therefore a function of the physicochemical properties of the proteins, nanoparticles, and surrounding fluid.^3,4^ This adsorbed phase displayed on the nanoparticle surface greatly impacts the way in which the nanoparticle interacts with its environment, dictating function, immunogenicity, localization, toxicity, and uptake.^5,6^ However, because the structure of the existing nanoparticle corona is often difficult to study and predict directly, despite theoretical progress to date^9^, the empirical study of solution phase protein-nanoparticle interactions is critically important, with a dearth of such information in the literature currently.^7–9^ Understanding protein-corona interactions is also central to their control in the context of nanoparticle sensors or nanosensors^24-26^. In this work, we address this problem by systematically examining the adsorption of 20 proteins to a well-studied carbon nanotube corona phase (CP), single-stranded (ss) (GT)_15_-DNA wrapped SWCNT (G-SWCNT). Our results provide new insights into the nature of binding and photophysical changes that occur to the latter, promising new sensor and probe design based on these findings.

Many fluorescent nanoparticles exhibit sensitivity to binding at the particle surface, with photophysical properties especially sensitive to protein adsorption, forming the basis of novel sensor technologies^10^. For example, these changes in nanoparticle properties resulting from the protein corona have been leveraged across a broad range of applications, including protein fingerprinting diagnostics and corona-mediated sensing approaches^10^, as well as improved therapeutic efficacy^6^, cellular uptake, and biocompatibility^11–14^ of nanoparticle-based medicines. Our laboratory has utilized engineered corona phases to control selective molecular binding at the carbon nanotube surface using a technique we refer to as Corona Phase Molecular Recognition (CoPhMoRe).^15–19^ The ability to predict how protein adsorption and complexation with the nanoparticle corona proceeds mechanistically, and the corresponding change in photophysical properties, would greatly accelerate innovation in this area.^20^

Fluorescent nanoparticles are particularly useful for monitoring protein adsorption and corona formation, since their photophysical properties vary strongly as a function of their local surface environment, enabling *in situ* characterization.^21^ Commonly, changes in ensemble fluorescence properties such as emission intensity, wavelength, and lifetime have been used to track protein adsorption.^22^ In addition, several techniques have been developed to yield more detailed information on protein-nanoparticle interactions. For instance, fluorescence correlation spectroscopy (FCS) has been used to track changes in nanoparticle size upon protein binding and to quantify nanoparticle-protein binding affinities.^23,24^ In this approach, fluorescence intensity fluctuations of particles are tracked as they diffuse in and out of a small focal volume. The temporal autocorrelation function of the intensity time trace can be related back to a hydrodynamic radius, which can be fit to a Hill equation as a function of protein concentration to yield a dissociation constant. Fluorescence resonance energy transfer (FRET) has also been employed to monitor protein adsorption using dye-labeled albumin with high spectral overlap with quantum dots.^24,25^ Here, the FRET efficiency as a function of protein concentration was similarly fit to a Hill equation to describe protein affinity. Emission from fluorophore-labeled polystyrene nanoparticles has also been used to track nanoparticle movement against separately labeled proteins to characterize the evolution of hard (strongly adsorbed) and soft (weakly adsorbed) CP formation.^26^

In the context of SWCNTs, photoluminescence (PL) changes arising from protein adsorption have been broadly applied to develop sensing constructs and characterize corona formation in complex media.^27–31^ When singly dispersed, semiconducting SWCNTs emit band-gap fluorescence in the near-infrared (nIR) and exhibit pronounced sensitivity to changes in their local dielectric and chemical environment, registered as changes in PL intensity (ΔI/I_0_) and emission wavelength (Δλ).^32–35^ This sensitivity has been leveraged for specific protein-sensing constructs, in which proteins have been interfaced with SWCNTs through protein adsorption, covalent conjugation, or recognition elements, including natural binders such as antibodies and aptamers, as well as synthetic CPs formed by the CoPhMoRe method.^15–19^ In these nanosensor constructs, changes in SWCNT PL serve as a quantitative readout of the presence of specific proteins or their activity. An active area of research involves understanding how competitively adsorbing proteins complex with, modify or displace coronas such as those from DNA oligonucleotides on SWCNT.^20^ Our current study is distinct from past efforts in its examination of a large slate of 20 proteins, systematically studying their solution-phase adsorption in a way that may form the basis for future predictive tools for nanosensor design.

Herein, we systematically evaluate protein adsorption to G-SWCNT using a panel of 20 proteins spanning a broad range of physicochemical properties. We quantify both PL intensity changes and wavelength shifts as a function of protein adsorption and relate these responses to effective protein binding affinities. We confirm that intensity and wavelength shifts can be complimentary readouts of protein interactions and that 15 of the 20 proteins follow a Langmuir binding isotherm. A minority of proteins (5) exhibit a non-monotonic two-step response in which an additional adsorption event at higher protein concentrations attenuates quenching or produces a net turn-on signal. Notably, we find that metalloproteins, proteins which contain metal ion cofactors, tend to produce larger quenching responses to protein adsorption compared to non-metalloproteins, providing a basis for increasing nanosensor signal generation. We advance a model where quenching is inversely correlated with protein size, with smaller proteins inducing greater quenching, suggesting a distance-dependent interaction between the metal cofactor and the SWCNT surface. Using horseradish peroxidase (HRP) as an example, we show that HRP apoprotein (apo-HRP) induces much less PL quenching compared to holo-HRP, highlighting the key role of the metal cofactor in modulating SWCNT PL on G-SWCNT. Together, these findings quantitatively describe how adsorption of diverse proteins can be detected on nanosensors like G-SWCNT and reveal an advantage that inorganic moieties can provide in potentially increasing photophysical transduction. This work in turn should be valuable for providing input for future predictive tools that allow the design of nanosensor corona for targeting specific proteins, a longstanding goal of the field.

## Results and Discussion

### PL Responses of SWCNTs across a Diverse Protein Panel

Singly dispersed SWCNTs have been developed for applications in a wide range of biological environments, including detection of diabetes and inflammation biomarkers in humans^30,31^, oxidative stress monitoring in plants^36–38^, and real-time tracking of reproductive markers in marine animals.^39^ In each of these settings, nanotubes encounter complex protein mixtures that can alter their photophysical properties and influence downstream function. To reflect the functional diversity of proteins present in such media, we screened a panel of 20 proteins spanning a broad range of physicochemical properties and conducted a systematic evaluation of how protein adsorption perturbs SWCNT photophysics. The panel included proteins of diverse biological origin (animal, plant, fungus) and molecular weight (8.5-580 kDa). All proteins were tested against the G-SWCNT construct in MES pH 5.7 buffer. We selected G-SWCNT for two key reasons: (i) it is one of the most widely studied constructs, enabling comparison with prior protein– SWCNT interaction studies^20^; and (ii) it is routinely deployed as a sensor in both plant and animal systems, making it an especially relevant platform for assessing how diverse proteins associate with and modulate this commonly used CP.^38,40^ MES was selected to match the buffer conditions routinely used for plant-associated experiments, and it is noteworthy that at this pH, the SWCNT is partially protonated.^41^

Proteins in solution were titrated into G-SWCNT suspensions at concentrations ranging from 10^-6^ mg/mL to 1 mg/mL, and the nIR spectrum was recorded before and after each addition (**Fig. 1a**). Proteins modulate SWCNT emission in two orthogonal ways: (i) intensity change (ΔI/I_0_), reflecting altered radiative versus non-radiative exciton decay pathways;^42,43^ and (ii) wavelength shift (Δλ), a solvatochromic response to local dielectric rearrangement and reorganization of the corona.^19,33,44^ Although SWCNTs synthesized using the High-Pressure Carbon Monoxide (HiPCO) technique contain multiple chiralities, the (6,5) chirality (emission peak at ∼990nm) dominates in our sample and was thus used for quantifying ΔI/I_0_ and Δλ. Because both ΔI/I_0_ and Δλ arise from protein–CP interactions at the nanotube surface, the dose–response behavior is expected to scale with the fraction of SWCNT corona adsorption sites occupied at equilibrium. As such, we modeled the dose-dependent changes in PL intensity and wavelength using a first-order reversible binding interaction between the protein (P) and the available adsorption sites (θ) on the G-SWCNT construct:

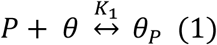

**Figure 1.**
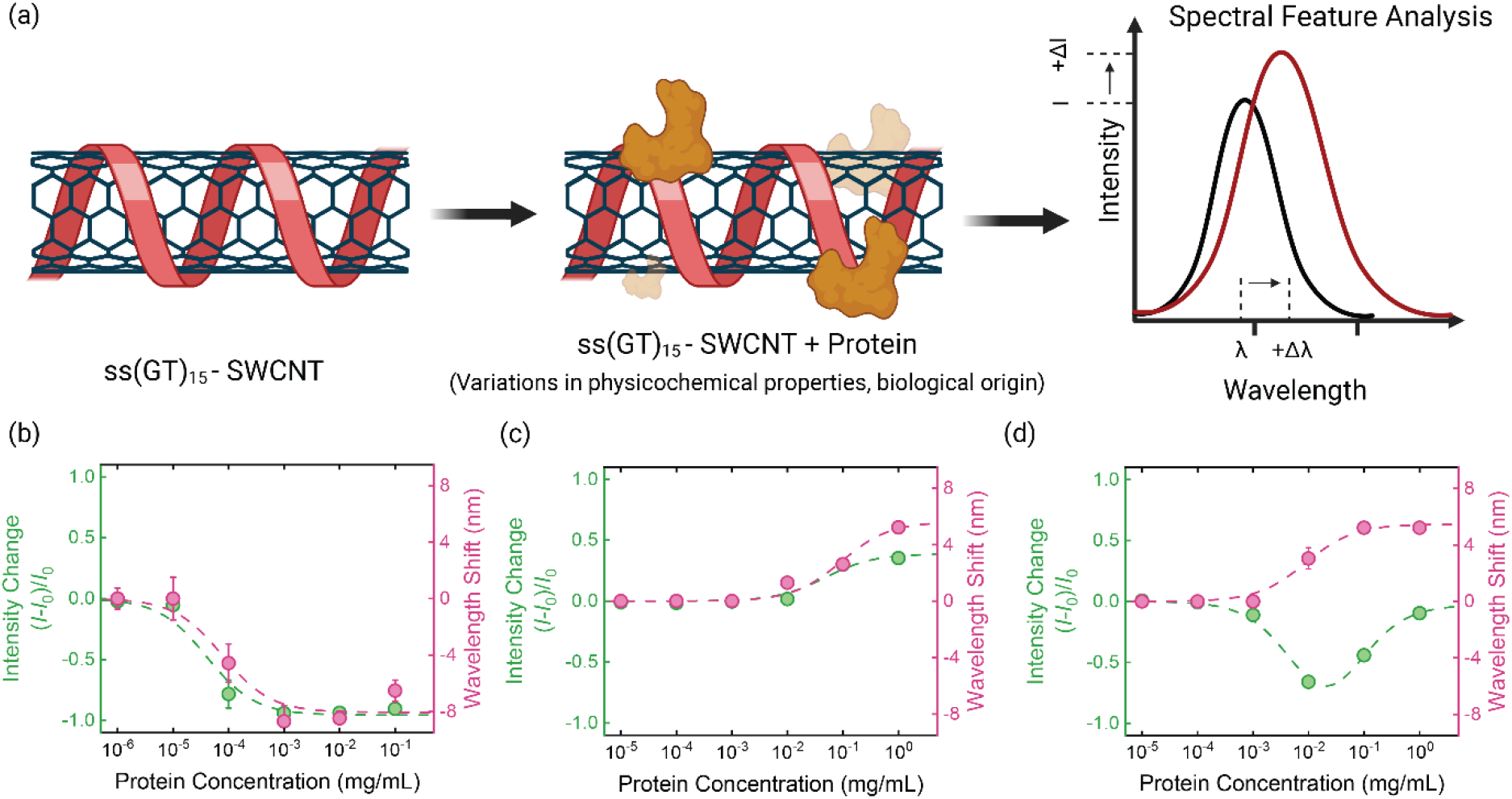
Protein binding alters the optical signature of ssDNA-wrapped SWCNTs. (a) Experimental workflow: G-SWCNT are incubated with individual proteins that differ in physicochemical properties and biological origin; adsorption perturbs the near-infrared emission spectrum, altering intensity (*ΔI*) and wavelength (Δ*λ*). (b–d) Representative Langmuir binding isotherms for three proteins: (b) Horseradish peroxidase (HRP); (c) alcohol dehydrogenase ADH; (d) arginase. Data points show measured (*I-I*_0_*)/I*_0_ (green, left axis) and Δλ (pink, right axis) as a function of protein concentration, with dashed curves indicating model fits that yield *K*_D,I_ and *K*_D,λ_.

The equilibrium for this reaction is 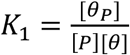. We assume that the fluorescence response is proportional to the fraction of occupied sites 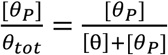, yielding a single-site Langmuir isotherm description of the observed spectral changes^45^:

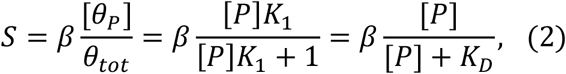

where *S* is the measured signal in terms of fluorescence intensity change 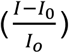 or wavelength shift Δλ = (λ − λ_0_); [*P*] is the protein concentration; 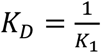is the dissociation constant in the same units as P; and β scales the magnitude of the photophysical response. All signals were background-corrected prior to fitting by subtracting the response of a matched buffer-only control.

**Figures 1b & c** illustrate representative fits for two proteins: horseradish peroxidase (HRP, 44 kDa), which yields near-complete quenching accompanied by a ∼9 nm blue-shift at higher concentrations, and alcohol dehydrogenase (ADH, 81 kDa), which produces a turn-on fluorescence response and a ∼5 nm red-shift at 1 mg/mL. Across the full panel of 20 proteins tested, we observed substantial differences in photophysical behavior. 15 proteins elicited both optical readouts (intensity modulation and wavelength shift), three proteins produced exclusively one type of response, and two proteins (chitinase & pectinase) induced no measurable signal change under our experimental conditions. These trends underscore that ΔI/I_0_ and Δλ report on distinct yet sometimes overlapping mechanisms. Because these processes can occur independently or concurrently, we extracted separate dissociation constants, *K*_*D,I*_ and *K*_*D*,λ_ , for each protein. The set of dissociation constants obtained from the single-site Langmuir model is reported in **Table 1**, and the corresponding fits are provided in **Supporting Figures 1-2**.

**Table 1.** Protein panel and single-site Langmuir fit parameters for adsorption to G-SWCNT.

| Protein: | PDB ID | Size (kDa) | $K_{D,I}$ (μM) | $K_{D,\lambda}$ (μM) | Cofactor | $\Delta I_{Max}/I_0$ | $\Delta\lambda_{Max}$ (nm) |
| --- | --- | --- | --- | --- | --- | --- | --- |
| Ubiquitin | 1UBQ | 8.5 | / | 1.8 | / | -0.08 | 5 |
| Actin | 3HBT | 42 | 0.18 | 0.81 | / | 0.17 | 6 |
| Subtilisin | 1SBC | 30 | / | 4.8 | / | 0.02 | 3 |
| Pectinase | 1BHE | 43 | / | / | / | 0.12 | 0 |
| Calmodulin | 1CLL | 19 | 5.6 | 1.0 | / | 0.34 | 4 |
| Chitinase | 1CTN | 30 | / | / | / | -0.05 | 0 |
| Alcohol Dehydrogenase (ADH) | 1HTB | 81 | 0.59 | 1.2 | Zn | 0.35 | 5 |
| Superoxide Dismutase (SOD) | 1Q0E | 32.5 | 0.19 | 2.1 | Cu, Zn | -0.50 | 2 |
| Urease | 7KNS | 582 | 4.8E-3 | 0.14 | Ni | -0.29 | 4 |
| Enolase | 1IYX | 93 | 0.1 | 0.21 | Mg | -0.19 | 4 |
| Ferritin | 6WX6 | 440 | 2.7E-3 | 0.11 | Fe | 0.26 | 4 |
| Horseradish Peroxidase (HRP) | 1HCH | 44 | 9.2E-4 | 1.8E-3 | Fe | -0.94 | -9 |
| RuBisCO | 1RCX | 550 | / | 0.037 | Mg | 0.01 | 3 |
| Catalase | 1TGU | 243 | 0.25 | 0.12 | Fe | -0.16 | 5 |
| Laccase | 1GYC | 70 | 0.59 | 1.1 | Cu | -0.86 | -8 |

PDB = protein data bank

While the single-site Langmuir model captured the dose–response behavior of most proteins, a subset of proteins displayed distinct signatures. Specifically, arginase, cytochrome c, hemoglobin, human serum albumin, and bovine serum albumin exhibited non-monotonic responses in ΔI/I_0_ and/or Δλ. For instance, PL intensity decreased at low protein concentrations and then increased at higher concentrations for arginase (**Fig. 1d**). These signatures cannot be captured by a single-site Langmuir model, which assumes monolayer coverage and non-interacting adsorbates. Instead, the data can be described by a two-step binding mechanism that allows for protein-protein interaction or intra-protein changes on adsorption. In this model, a protein first binds to an unoccupied SWCNT site to produce a turn-off response; at higher concentrations, the protein additionally interacts with a pre-bound site to generate an opposing turn-on (or weaker turn-off) response. This two-step binding mechanism has previously been applied to describe similar non-monotonic PL changes in an interleukin-6 SWCNT sensor^29^ and separately a SWCNT nanosensor for fibrinogen.^19^ The response in these cases could also result from a parallel two step mechanism in which proteins at high concentration first associate in solution, and the resulting complex adsorbs to the SWCNT, leading to the same final state.

For these cases, we describe a reversible two-step adsorption model assuming two equilibria:

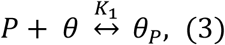

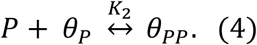

The resulting total site balance is

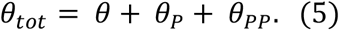

At equilibrium,

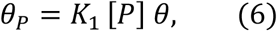

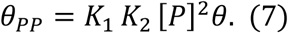

The fluorescence response can then be written as

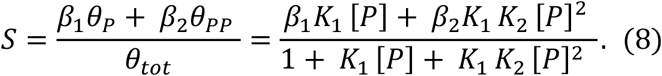

Here, *K*_1_ and *K*_2_ represent the binding affinities for the first and second adsorption steps, respectively. Because the PL intensity decrease occurs at low concentrations and the turn-on (or weaker quenching) response emerges only at higher concentrations, the characteristic transition concentration satisfies 1/K_1_ < 1/K_2_, consistent with a stronger first interaction. The parameters β_1_ and β_2_ scale the magnitude and sign of the photophysical response associated with each bound state. In all proteins exhibiting this two-stage behavior, the second state θ_*PP*_ resulted in either overall less quenching (0 > β_2_ > β_1_) or a turn-on response (β_2_ > 0). Note that an equivalent equilibrium form is obtained in the alternative mechanism in which proteins first associate in solution (P + P ⇌ PP) before adsorbing to SWCNT (**Supporting Equations S1-S4**). Thus, this two-step model can be interpreted as capturing either surface-assisted binding or solution-phase association followed by adsorption.

In applying this model to the experimental data, we observed that the two-step mechanism manifested differently across the two optical channels. For two of the five proteins, non-monotonic behavior appeared in both ΔI/I_0_ and Δλ, indicating that the second adsorption event not only altered exciton quenching pathways but also reorganized the local dielectric environment. For the other proteins, however, the two-step signature manifested only in intensity. This may suggest that protein-protein assemblies formed at higher concentrations can modulate exciton decay pathways differently from individually adsorbed proteins, without producing corresponding non-monotonic changes in emission wavelength indicative of additional corona reorganization. For proteins showing no evidence of such phenomena in the wavelength channel, the single-site Langmuir model introduced above was applied. Note that setting *K*_2_ = 0 in **Equation 8** recovers the functional form of the single-site adsorption model from **Equation 2. Table 2** summarizes the extracted dissociation constants for both optical channels for proteins following the two-step mechanism and **Supporting Figures 1-2** provide the full set of fitted calibration curves. Notably, BSA exhibited approximately equal apparent binding affinities for both steps (1/K_1_ = 1/K_2_ ≈ 0.25 µM), which may indicate that the two steps occur over similar concentration scales and manifest in the optical response as a coupled transition.

**Table 2.** Protein subset exhibiting non-monotonic responses with two-step adsorption model fit parameters for adsorption to G-SWCNT.

| Protein: | PDB ID | Size (kDa) | $1/K_{1,I}$ ($\mu\text{M}$) | $1/K_{2,I}$ ($\mu\text{M}$) | $1/K_{1,\lambda}$ ($\mu\text{M}$) | $1/K_{2,\lambda}$ ($\mu\text{M}$) | Cofactor | $\Delta I_{\text{Max}}/I_0$ | $\Delta\lambda_{\text{Max}}$ (nm) |
| --- | --- | --- | --- | --- | --- | --- | --- | --- | --- |
| Bovine Serum Albumin (BSA) | 9QQD | 66 | 0.25 | 0.25 | 0.49 | / | / | -0.20 | 4 |
| Human Serum Albumin (HSA) | 1E7I | 66.5 | 0.089 | 4.3 | 0.25 | / | / | -0.12 | 5 |
| Hemoglobin | 2HHB | 64.5 | 0.073 | 1.4 | 0.084 | 0.30 | Fe | -0.73 | -5 |
| Cytochrome C (CTC) | 1HRC | 12.4 | 0.072 | 0.44 | 0.11 | 1.6 | Fe | -0.73 | -5 |
| Arginase | 1RLA | 120 | 0.050 | 0.56 | 0.074 | / | Mn | -0.66 | 5 |

These dissociation constants describe protein adsorption to the engineered corona phase (G-SWCNT) and should be characterized as apparent binding affinities measured after both protein binding and photophysical interaction with G-SWCNT occur. Further, this adsorption process represents the formation of a biocorona, defined here as a subset of the term corona phase, in which proteins and other biomolecules spontaneously associate with the particle surface from biological media via non-covalent interactions.^6,46^ Tabulating these dissociation constants therefore allows for comparisons of protein affinities and provides training data for computational models that seek to predict SWCNT–protein interactions from molecular features. Across proteins for which both *K*_*D,I*_ and *K*_*D*,λ_ were obtained, the two dissociation constants typically fell within one order of magnitude of each other, reflected by their proximity to the parity line in log–log space (**Supporting Figure 4**). A modest trend toward *K*_*D,I*_ < *K*_*D*,λ_ was observed, suggesting that proteins may influence exciton quenching pathways at slightly lower surface coverage than is required to measurably perturb the local dielectric environment. These relationships offer a useful reference for comparing the relative sensitivities of intensity and wavelength changes due to protein adsorption for this CP (G-SWCNT).

### Most Metalloproteins Quench G-SWCNT More than Non-Metalloproteins

The PL intensity changes observed across our protein screen included both turn-on and turn-off responses, indicating that protein adsorption can enhance radiative exciton recombination, either directly or through interacting with the DNA, or introduce additional non-radiative decay pathways. In practice, because SWCNT PL depends strongly on the local electronic environment, adsorption of a molecule that places quenching motifs near the nanotube surface can modulate PL intensity by altering non-radiative decay pathways and/or the occupancy of electronic states involved in exciton formation. For example, pronounced turn-off responses often arise when metal ions are brought into proximity to the SWCNT surface through charge or energy transfer interactions.^47,48^ Many proteins themselves contain bound metal ions or metal-containing clusters (for example, Fe, Cu, or Zn) that support key biological functions including electron transfer, oxygen transport, and catalytic redox chemistry.^49^ Proteins that incorporate such metal cofactors are broadly classified as metalloproteins, and roughly half of all proteins fall into this category. With this in mind, we grouped the protein panel by cofactor content. Here, 12 of the 20 proteins contained metal cofactors, allowing us to test whether metalloproteins exhibit different PL modulation than metal-free proteins.

**Figure 2a** summarizes the extent to which each protein modulates G-SWCNT PL intensity, and **Figure 2b** provides representative nIR spectra showing the concentration-dependent PL response for HRP as an example. Because several proteins exhibited non-monotonic dose–response behavior, we used the maximum intensity change observed across the titration series for each protein as a standardized metric. Differentiating proteins by cofactor content revealed a clear distinction in their photophysical effects. Metalloproteins tended to produce greater quenching responses than metal-free proteins, with maximum ΔI/I_0_ values of 94% for metalloproteins versus 20% for metal-free proteins. This trend is consistent with the expected ability of metal centers to introduce additional non-radiative exciton decay pathways.^47,48,50–53^ However, the metalloprotein cohort also displayed some heterogeneity, with several proteins producing comparatively weak quenching and even some turn-on responses. In particular, alcohol dehydrogenase (ADH), a Zn-containing metalloprotein, produced the largest turn-on response within the metalloprotein cohort.

**Figure 2.**
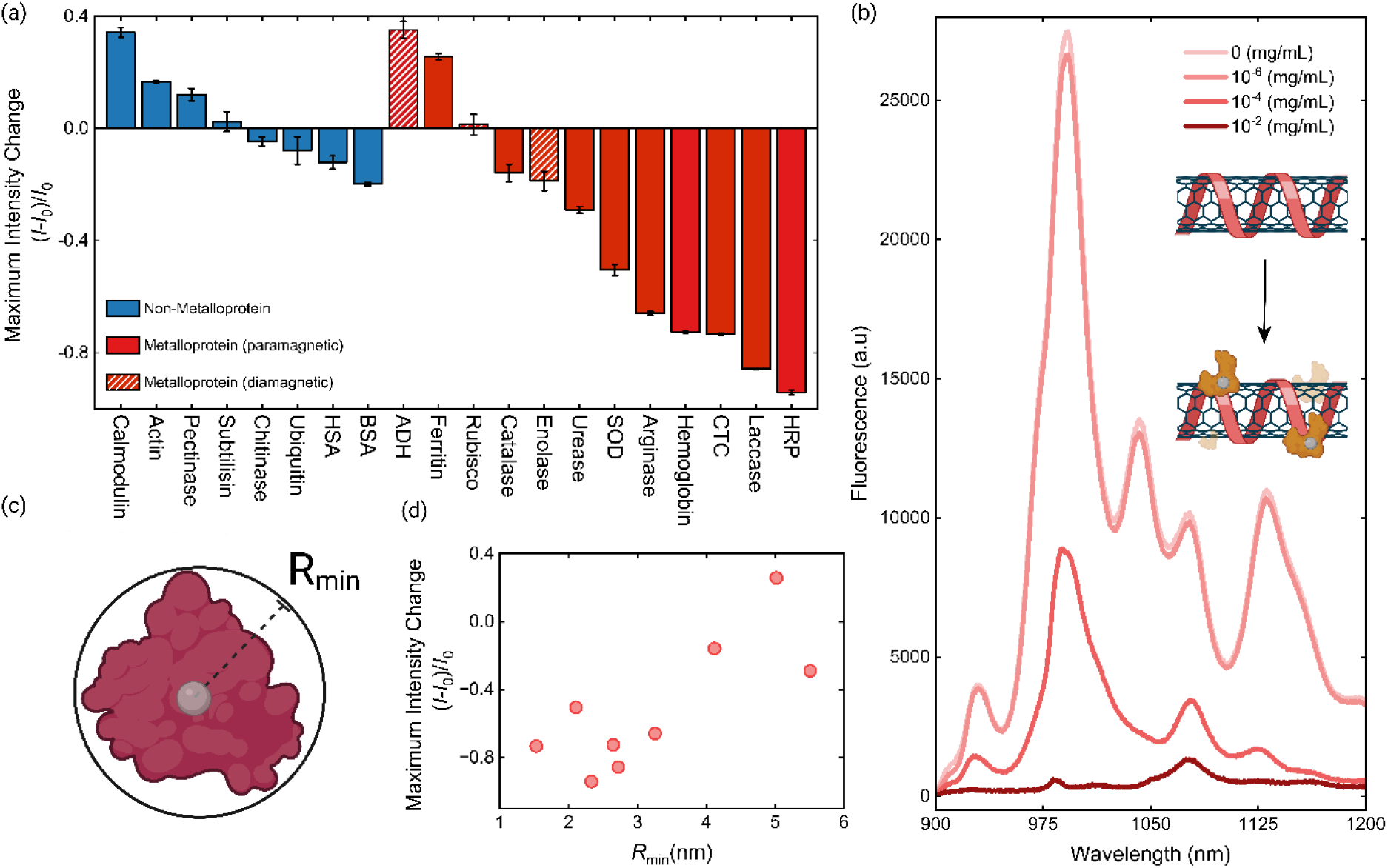
Metal cofactors tend to increase photoluminescence quenching of G-SWCNT on protein adsorption. (a) Maximum normalized intensity changes for each protein, color-coded by class (blue = non-metal proteins, red = metalloproteins. red with dashed line = diamagnetic metal cofactor). (b) Representative NIR emission spectra of G-SWCNT before and after titration with HRP. (c) Schematic representation of the minimum spherical radius (*R*_min_) of a protein, used as a proxy for metal-to-SWCNT separation. (d) Maximum intensity change plotted against R_min_ for paramagnetic ion containing metalloproteins (r = 0.777).

This spread in metalloprotein responses motivated us to examine whether metal cofactor identity, rather than metalloprotein classification alone, contributes to quenching strength. Notably, among the weaker-quenching metalloproteins, several contain diamagnetic cofactors (Mg^2+^ or Zn^2+^). Accordingly, we separated metalloproteins by whether their cofactors were paramagnetic or diamagnetic. Of the twelve metalloproteins in our screen, nine contain paramagnetic cofactors (Fe^+3^/Fe^2+^, Cu^2+^, Mn^2+^, Ni^2+^), while the remaining three (ADH, enolase, and RuBisCO) contain only diamagnetic ions (Mg^2+^ or Zn^2+^). Separating these two classes revealed that paramagnetic metalloproteins quenched G-SWCNT emission more strongly than their diamagnetic counterparts, with maximum ΔI/I_0_ quenching values of 94% compared to 19% for the diamagnetic metalloproteins.

A similar trend has been reported in earlier ion-screen studies using sodium dodecylbenzene sulfonate (SDBS) wrapped SWCNTs, where paramagnetic metal ion salts were found to quench the nIR SWCNT emission more than diamagnetic ions.^48^ Because those experiments involved free ions rather than proteins, the enhanced quenching could be more directly attributed to the ion–nanotube interaction itself. Notably, although the paramagnetic ions quenched more strongly, the quenching efficiency did not correlate with magnetic moment, indicating that paramagnetism was not the dominant mechanism. Instead, the authors attributed the increased quenching to short-range electronic interactions arising from high local ion concentrations at the nanotube surface, suggesting that ion proximity to the SWCNT, rather than paramagnetism, was the more significant determinant of quenching degree. In our system, metal ions are protein-bound and interact with DNA-wrapped SWCNTs through a complex CP. Consequently, while factors such as cofactor accessibility and proximity to the SWCNT surface likely influence PL quenching, the limited number of exclusively diamagnetic metalloproteins in our dataset and the complexity of the protein-SWCNT interaction preclude a more detailed mechanistic distinction.

One outlier, Ferritin, contains iron but produced a moderate turn-on response. Ferritin is structurally distinct; as an iron storage protein, it comprises an approximately spherical 24-subunit assembly that sequesters thousands of Fe^3+^ ions within a thick (∼2 nm) protein shell.^54^ As a result, the metal cofactor in ferritin may be sterically blocked from the SWCNT surface even after protein adsorption.

To probe the role of proximity more directly, we treated each protein as a simple sphere, assuming the most compact geometry possible, and converted its molecular weight to a minimum radius with:

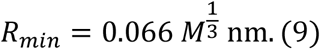

where *R*_*min*_ is the radius in nm of the smallest sphere that could contain the given protein mass and *M* is the molecular weight in Da (**Fig. 2c**).^55^ This first-order metric serves as an approximate proxy for metal–SWCNT separation and therefore allows us to test whether proximity of a metal cofactor corresponds to stronger quenching. Focusing this analysis on the paramagnetic metalloprotein subset, which showed the greatest quenching responses, we observed an inverse trend between Δ*I*/*I*_0_ and *R*_min_: smaller proteins generally induced greater quenching (r = 0.777, **Fig. 2d**). While informative, this model provides only a crude approximation of the cofactor–nanotube separation. It assumes that the metal center resides at the geometric center of a rigid spherical protein, which is unlikely to hold across all metalloproteins. In practice, proteins may adopt conformations that can either shield the metal cofactor from the nanotube surface or, conversely, expose it more directly upon adsorption.^56^ In some cases, proteins may also denature upon contact with the SWCNT surface.^57–59^ More refined structural metrics, such as nanotube docking geometries, would likely strengthen the observed correlations. Nevertheless, this coarse approximation suggests that the proximity of a metal center to the SWCNT surface is an important determinant of the quenching magnitude.

Taken together, these results suggest that metal–SWCNT separation contributes to cofactor-mediated quenching. Proximity-dependent metal quenching has also been intentionally exploited in engineered SWCNT sensing constructs, providing a useful point of comparison. For example, this interpretation is consistent with our prior work on immunoglobulin G (IgG1) detection, in which we used a Cu(II)/protein A complex to pre-quench SWCNT fluorescence and observed a turn-on response upon IgG1 binding, attributed to an increase in the Cu(II)–nanotube separation.^60^ More recently, Ledesma et al. used covalent attachment of HRP to SWCNTs for selective sensing of its native substrate, H_2_O_2_, by leveraging quenching from the heme cofactor positioned near the nanotube surface.^61^ In contrast to these engineered protein-SWCNT constructs, our results show that the same proximity-dependent modulation of SWCNT PL can emerge from protein adsorption itself.

To distinguish metal cofactor-mediated PL quenching from alternative mechanisms, we evaluated two competing mechanisms, cofactor-independent protein binding to SWCNTs and protein-induced SWCNT aggregation. In the former mechanism, equivalent to standard CoPhMoRe, specific interactions between proteins and the SWCNT CP modulate PL intensity independently of metal cofactors. In the case of SWCNT aggregation, inter-tube interactions within bundles introduce nonradiative decay pathways that quench PL.^34^ We examined HRP as a model system, comparing SWCNT responses to its holo- and apoprotein forms (**Fig. 3a**). HRP was selected because it is well characterized, and apo-HRP is commercially available, enabling a direct comparison in the presence and absence of the heme prosthetic group in HRP. Importantly, apo-HRP largely maintains its secondary structure under native buffer conditions, making it a suitable comparator for isolating the effect of the metal cofactor.^62,63^

**Figure 3.**
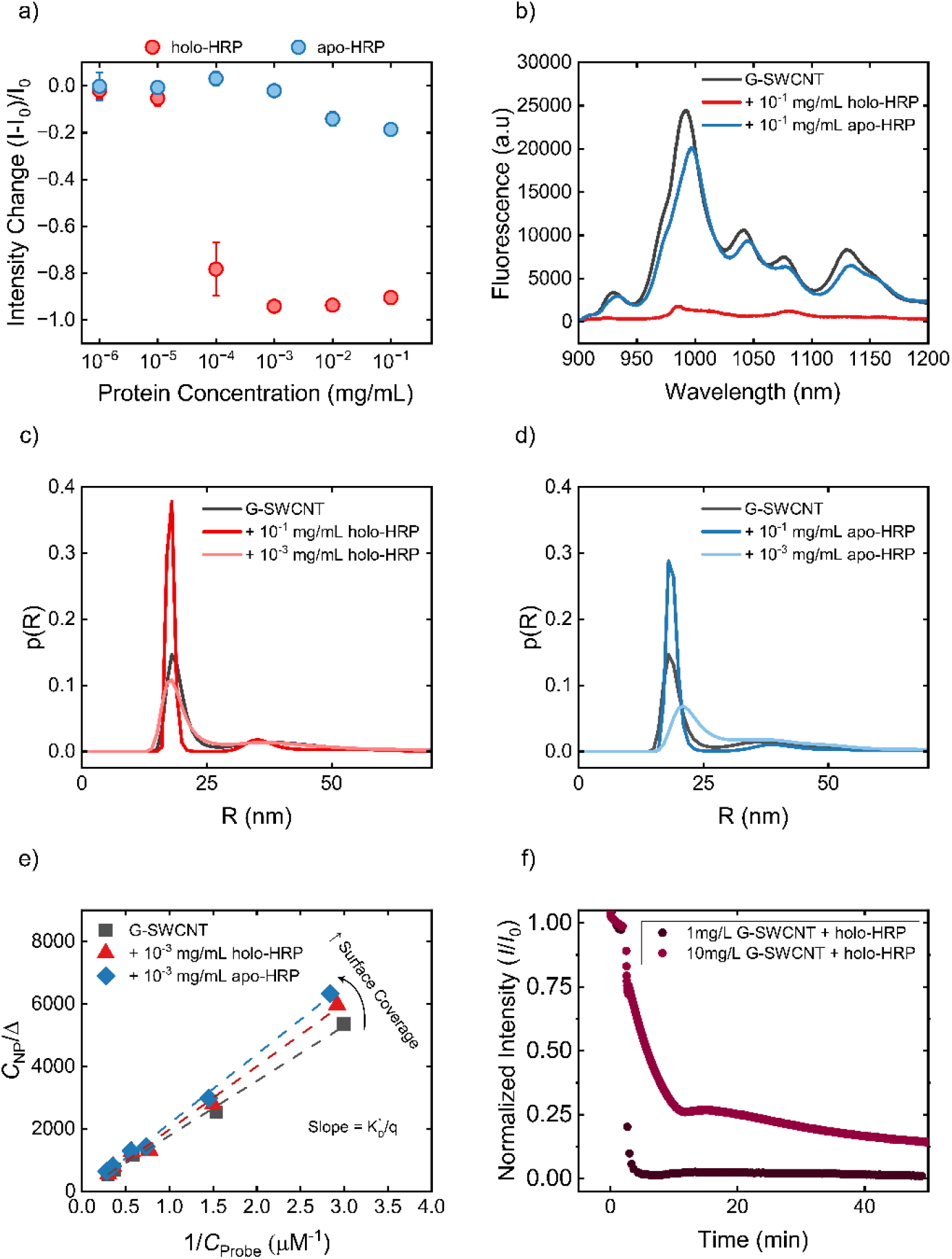
Metal cofactors are required for PL Quenching. (a) PL intensity change ((*I-I*_0_*)/I*_0_) as a function of protein concentration for holo-HRP (red, contains heme) vs. apo-HRP (blue, heme removed). Holo-HRP fully quenches SWCNT PL (ΔI/I_0_ = −0.94), whereas apo-HRP only moderately quenches PL intensity (ΔI/I_0_ = −0.19). (b) Representative nIR emission spectra of G-SWCNT before and after mixing with apo- or holo-HRP at 0.1 mg/mL and incubating for 1 hour. (c-d) Hydrodynamic radius determined by single particle tracking analyzed with MApNTA for G-SWCNT alone or mixed with (c) holo-HRP or (d) apo-HRP. Incubation with 10^-3^ mg/mL protein shows broadened size distributions consistent with adsorption for both cases. (e) Molecular probe adsorption (MPA) analysis indicates reduced accessible surface area on the SWCNT surface 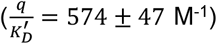 after incubation with apo-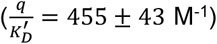 or holo-HRP 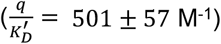 at 10^-3^ mg/mL, consistent with adsorption. (f) PL Intensity vs. time for 1 mg/mL or 10 mg/mL SWCNTs incubated with 10^-3^ mg/mL holo-HRP. Slower quenching kinetics with higher SWCNT concentration are inconsistent with a protein-induced SWCNT aggregation-based mechanism.

As in the previously described protein screen, apo-HRP was titrated into G-SWCNT suspensions at concentrations ranging from 10^-6^ mg/mL to 10^-1^ mg/mL, and the nIR spectrum was recorded before and after each addition (**Fig. 3b**). Compared to holo-HRP, apo-HRP induced significantly less quenching, with a Δ*I*_*Max*_ of -0.19 vs. -0.94. Interestingly, both *K*_*D,I*_ and *K*_*D*,λ_ were substantially higher for apo-HRP (1.2 × 10^-7^ M and 1.7 × 10^-7^ M, respectively) compared with holo-HRP (9.2 × 10^-10^ M and 1.8 × 10^-9^ M). The difference in *K*_*D,I*_ may in part reflect the smaller intensity response observed for apo-HRP, limiting the robustness of model fitting. However, the higher *K*_*D*,λ_ for apo-HRP is unexpected, given that apo-HRP is reported to have a similar overall structure to holo-HRP and would therefore be anticipated to induce comparable adsorption-driven rearrangements of the SWCNT corona.

To confirm adsorption of apo-HRP at similar concentrations to holo-HRP, we performed single particle tracking with maximum *a posteriori* nanoparticle tracking analysis (MApNTA) to measure changes in hydrodynamic radius after incubating G-SWCNT with HRP (**Fig. 3c-d**).^64^ We measured G-SWCNT alone, and G-SWCNT incubated with the upper (10^-1^ mg/mL) and lower (10^-3^ mg/mL) concentrations of HRP that yielded full PL quenching for holoprotein. At 10^-3^ mg/mL protein, both apo- and holo-HRP showed broadened size distributions consistent with adsorption, with apo-HRP additionally showing an increased mode. Surprisingly, at 10^-1^ mg/mL protein, the size distributions narrow but retain the same mode as G-SWCNT alone. We hypothesize that this could be due to protein-protein interactions affecting binding to the SWCNT surface, as described earlier in the two-step adsorption model for non-monotonic behavior. Additionally, at 10^-1^ mg/mL protein, the measured particle number density (∼7.5 × 10^9^ particles/mL) exceeded the recommended range for single particle tracking (≤10^9^ particles/mL), which may affect the reliability of the inferred size distributions.

We further confirmed protein adsorption at 10^-3^ mg/mL using molecular probe adsorption (MPA).^65,66^ In this technique, riboflavin functions as a reporter of solvent-exposed SWCNT surface area and is quenched upon surface adsorption. After incubating varying concentrations of riboflavin with SWCNT, the remaining unadsorbed riboflavin can be compared to the total amount added to infer relative SWCNT surface coverage from the governing equation

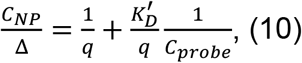

with Δ = *C* _*total*_ − *C* _*probe*_ . Here, *C* _*NP*_ is the SWCNT concentration, *C* _*total*_ is the total riboflavin concentration, *C* _*probe*_ is the unbound riboflavin concentration, *q* is the maximum amount of riboflavin that can adsorb per mass of SWCNT, and *K*^′^ is the dissociation constant of riboflavin to SWCNT. Because MPA infers surface coverage from the fraction of probe that is quenched, we additionally quantified and corrected for any non-SWCNT-related quenching arising from interactions of riboflavin with other solution components (e.g., free ssDNA and/or protein) by including a background correction term (*C* _*background*_); details of the background control and correction are provided in the Supplementary Information. The relative available SWCNT surface area is indicated by a value 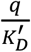, determined by the inverse of the slope of 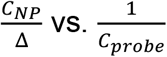, where a smaller value represents less available surface area. Compared to G-SWCNT alone 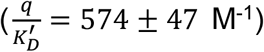, G-SWCNT incubated with holo-HRP 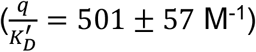 or apo-HRP 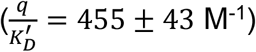 at 10^-3^ mg/mL exhibited reduced accessible surface area (**Fig. 3e**), indicating protein adsorption. Further details on MPA are provided in the **Supporting Information**.

Together, MApNTA and MPA experiments suggest that the differences in PL responses from apo- and holo-HRP are not due to major differences in adsorption affinity.

Finally, to investigate the possibility of protein-induced SWCNT aggregation, we analyzed the kinetics of SWCNT PL decay upon introduction of holo-HRP (**Fig. 3f**). In an aggregation-based mechanism, PL quenching should occur faster with higher SWCNT concentration at fixed HRP concentration due to increased SWCNT collision frequency. We observe the opposite trend, with faster PL quenching at a lower SWCNT concentration. Consistent with these observations, the MApNTA measurements do not show large, multi-SWCNT aggregates in the size distributions. Overall, these complementary measurements demonstrate that HRP-induced PL quenching arises from interactions between SWCNT and the heme cofactor, rather than from cofactor-independent protein binding or protein-induced SWCNT aggregation.

### Conclusion

Nanoparticles hold great promise for probing molecular biology, yet their application is often complicated by protein adsorption. Such adsorption can alter nanoparticle properties and desired functions. In this work, we analyzed the photophysical responses of SWCNTs after protein adsorption using G-SWCNTs in MES buffer as a model system. We construct a dataset of 20 unique proteins varying widely in physicochemical properties and biological origin. We demonstrate how PL intensity and wavelength changes serve as orthogonal readouts of adsorption, reflecting protein-specific modulation of exciton relaxation pathways and nanoparticle corona structure. For 15 proteins, both intensity and emission wavelength changed monotonically with protein concentration and were well described by a one-step Langmuir adsorption model. The other five proteins varied non-monotonically in one or both PL readouts and were well described using a two-step model that additionally allows for protein-protein interactions that are likely to occur at higher concentrations.

The 12 metalloproteins in our dataset induced greater PL quenching compared to the non-metalloproteins, highlighting a role of metal cofactors in facilitating nonradiative exciton decay. This effect is further pronounced when considering only the nine paramagnetic ion-containing metalloproteins. The degree of quenching for the paramagnetic subset exhibits a size-dependence, with smaller proteins showing a higher degree of quenching. These findings are consistent with prior literature noting quenching effects of paramagnetic metal ions and distance-dependent metal sensing. We verify this metal-dependent mechanism by comparing SWCNT PL responses to holo-HRP and apo-HRP, which are structurally similar but differ in the presence or absence of the iron-containing heme group. Upon adsorption to SWCNT, the former induces near total quenching, whereas the latter is only modestly responsive in the intensity readout. We additionally rule out protein-induced SWCNT aggregation as a quenching mechanism by using single particle tracking analysis to estimate hydrodynamic radius and by comparing the SWCNT-concentration dependence of PL quenching kinetics. These results identify metalloproteins, especially those with paramagnetic metals, as a subset of proteins with an increased tendency to induce PL quenching in G-SWCNTs. Overall, we have developed a valuable dataset for understanding protein interactions with nanoparticles, quantifying PL changes across proteins spanning a broad range of physicochemical properties. The data should inform the design and interpretation of nanoparticle systems applied in biological settings.

## Materials and Methods

### Materials

Raw HiPCO SWCNTs were purchased from NanoIntegris (batch HR29-039). GT_15_ ssDNA oligonucleotides were obtained from Integrated DNA Technologies. 2-(N-morpholino)ethanesulfonic acid (MES) buffer was purchased from Fisher Scientific (cat. no. AAJ63341AK). For proteins, BSA, HSA (cat. no. A8763), hemoglobin (cat. no. H7379), CTC (cat. no. C2037), urease (cat. no. 666133), arginase (cat. no. A3233), ubiquitin (cat. no. U6253), ADH (cat. no. 74931), actin (cat. no. A2522), subtilisin (cat. no. P5380), pectinase (cat. no. P2401), calmodulin (cat. no. C4874), chitinase (cat. no. C6137), SOD (cat. no. S9697), enolase (cat. no. E6126), ferritin (cat. no. F4503), HRP (cat. no. P8375), apo-HRP (cat. no. P6278), RuBisCO (cat. no. R8000), catalase (cat. no. C40), and laccase (cat no. 38429) were purchased from Sigma Aldrich.

### Preparation of G-SWCNTs

A mixture of 2 mg ssDNA and 1 mg HiPCO SWCNTs was prepared in 1 mL of 100 mM NaCl in nuclease free water and sonicated using a 3 mm probe tip (Cole-Parmer) for 30 min at 40% amplitude in an ice bath (QSonica Q125). The resulting dispersion was centrifuged twice at 32,000 g for 90 min (Eppendorf 5430 R) to remove unsuspended SWCNT bundles. The concentration of the final G-SWCNT suspension was determined from its absorbance at 632 nm using an extinction coefficient of 0.036 mg L−1 cm−1 (Shimadzu UV-2600i). The sample was stored at 4 °C in a fridge for further use. G-SWCNT zeta potential was -30.47 ± 2.00 mV. Zeta potential was measured at 5 mg/L in 10 mM KCl + 10 mM Tris-EDTA buffer at pH 8.0 (NanoBrook 90Plus PALS, Brookhaven Instruments, see **Supplementary Information** for details).

### Protein Titrations and nIR Spectroscopy

High-throughput screening of protein adsorption onto the G-SWCNT construct was performed using a customized nIR microscope consisting of a Zeiss inverted microscope body with a 20× objective, coupled to an Acton SpectraPro 2500i spectrometer and a liquid-nitrogen-cooled InGaAs 1D detector (Princeton Instruments, 7514-0001). The G-SWCNT dispersion was diluted to 2 mg L−1 in MES buffer (pH 5.7). For titrations, 90 μL of G-SWCNT dispersion was combined with 10 μL of protein solution to achieve final protein concentrations spanning 10^-6^ mg/mL to 1 mg/mL in optically clear, flat-bottom 96-well plates (GenClone, 25-109). A matched buffer-only addition (90 μL G-SWCNT + 10 μL buffer) was used as a negative control. Plates were sealed with Parafilm and placed on a shaker plate to equilibrate at room temperature prior to measurement. Near-infrared fluorescence spectra were collected in triplicate from 890 to 1220 nm under 785 nm laser excitation (Invictus 785, Kaiser Optical Systems Inc., 365 mW). Spectra were processed to extract the (6,5) peak intensity and peak wavelength for each replicate, and the mean and standard deviation were reported for each condition.

### Model Fitting

Dose–response curves for both the normalized intensity change (ΔI/I_0_) and wavelength shift (Δλ) were fit in MATLAB using nonlinear least-squares optimization. Proteins exhibiting monotonic responses were fit to a single-site Langmuir isotherm using constrained nonlinear minimization (*fmincon*) with bounds enforcing *K*_*D*_. Proteins exhibiting non-monotonic two-stage responses were fit to the two-step adsorption model using *lsqcurvefit* with bounded parameters. For both models, β parameters were constrained to physically reasonable ranges to prevent implausible scaling and stabilize estimation of dissociation parameters, particularly for the two-step model. Specifically, intensity fits used β ∈ [− 1.1 , 1.1], and wavelength fits used β ∈ [− 10 , 10] nm.

### Kinetic measurements

Time-resolved measurements were performed by rapidly mixing horseradish peroxidase (HRP) with G-SWCNT suspensions directly in a 96-well plate and monitoring the near-infrared emission as a function of time using the same nIR microscope–spectrometer setup described above. Briefly, 10 μL of an HRP solution was added to 90 μL of G-SWCNT dispersion to yield a final HRP concentration of 1 μg/mL and final G-SWCNT concentrations of either 10 mg/L or 1 mg/L. Immediately after addition, samples were mixed by pipetting and spectral acquisition was initiated without delay. Sequential spectra were collected using exposure times of 2 s for the 10 mg/L G-SWCNT condition and 20 s for the 1 mg/L condition, and the (6,5) peak intensity was extracted for each time point to generate kinetic traces.

### Single Particle Tracking

A NanoSight LM10 (Malvern, USA) and NTA3.4 NanoSight software was used for single particle tracking of SWCNT. SWCNT were diluted to 1 mg/L in MES buffer (pH 5.7) and pumped through the imaging frame at 50 μL/min over 6s using a Harvard Apparatus (Holliston, MA) Pump 11 Elite syringe pump. For each sample, 16 videos were captured for 30s each. Temperature was held constant at 20 °C. Particle trajectories were analyzed with maximum *a posteriori* nanoparticle tracking analysis to estimate the hydrodynamic radius.

### Molecular Probe Adsorption

The MPA analysis method was performed as previously described. Briefly, SWCNT dispersions were diluted to 10 mg/L. For protein experiments, 10 μL of the protein solution was added to the SWCNT dispersions to achieve a final concentration of 10^-3 mg/mL. Samples were sealed with Parafilm and left to equilibrate on a shaker plate for 1 hour. Afterwards, riboflavin was dissolved in Nanopure water, and a stock solution was prepared at concentrations ranging from 0 to 50 μM. For each experiment, 10 μL of the riboflavin stock solution was added to the protein/SWCNT dispersions, resulting in a final concentration range of 0 to 5 μM. Fluorescence measurements were conducted by exciting riboflavin at 460 nm, with emission spectra collected from 480 to 600 nm in 2 nm increments using the top-reading position of a Varioskan Flash spectral scanning multimode microplate reader (Thermo Scientific).

## Acknowledgements

The authors are grateful for support from the Nanotechnology for Agricultural and Food Systems (A1511) program [grant no. 2021-67021-33999/project accession no. 1025638] under the USDA National Institute of Food and Agriculture. T. K. P. and G. S-V. are grateful for support from the National Science Foundation Graduate Research Fellowship Program under Grant No. 2141064.

## Notes

### Competing Interest Statement

The authors have declared no competing interest.

## References

1. Sun, Y., Zhou, Y., Rehman, M.Wang, Y.-F. & Guo, S. Protein Corona of Nanoparticles: Isolation and Analysis. Chem Bio Eng. 1, 757–772 (2024).

2. Cedervall, T. et al. Understanding the nanoparticle–protein corona using methods to quantify exchange rates and affinities of proteins for nanoparticles. Proc. Natl. Acad. Sci. 104, 2050–2055 (2007).

3. De, M., You, C.-C., Srivastava, S. & Rotello, V. M. Biomimetic Interactions of Proteins with Functionalized Nanoparticles: A Thermodynamic Study. J. Am. Chem. Soc. 129, 10747– 10753 (2007).

4. Tenzer, S. et al. Rapid formation of plasma protein corona critically affects nanoparticle pathophysiology. Nat. Nanotechnol. 8, 772–781 (2013).

5. Lee, S. et al. Corona Dynamics of Nanoparticles and Their Functional Design Space in Molecular Sensing. ACS Nano 19, 24653–24668 (2025).

6. Mahmoudi, M., Landry, M. P., Moore, A. & Coreas, R. The protein corona from nanomedicine to environmental science. Nat. Rev. Mater. 8, 422–438 (2023).

7. Vilanova, O. et al. Understanding the Kinetics of Protein–Nanoparticle Corona Formation. ACS Nano 10, 10842–10850 (2016).

8. Darabi Sahneh, F., Scoglio, C. & Riviere, J. Dynamics of Nanoparticle-Protein Corona Complex Formation: Analytical Results from Population Balance Equations. PLoS ONE 8, e64690 (2013).

9. Lundberg, D. J. & Strano, M. S. Approximate Corona Phase Hamiltonian for Individual Cylindrical Nanoparticle–Polymer Interactions. J. Phys. Chem. B 126, 347–354 (2022).

10. Cheeseman, S. et al. The Challenges and Opportunities of Protein Coronas for Nanoscale Biomolecular Sensing. Small 21, e03820 (2025).

11. Yin, H. et al. Reducing the cytotoxicity of ZnO nanoparticles by a pre-formed protein corona in a supplemented cell culture medium. RSC Adv. 5, 73963–73973 (2015).

12. Ritz, S. et al. Protein Corona of Nanoparticles: Distinct Proteins Regulate the Cellular Uptake. Biomacromolecules 16, 1311–1321 (2015).

13. Duan, G. et al. Protein corona mitigates the cytotoxicity of graphene oxide by reducing its physical interaction with cell membrane. Nanoscale 7, 15214–15224 (2015).

14. Corbo, C. et al. The Impact of Nanoparticle Protein Corona on Cytotoxicity, Immunotoxicity and Target Drug Delivery. Nanomed. 11, 81–100 (2016).

15. Mann, F. A., Herrmann, N., Opazo, F. & Kruss, S. Quantum Defects as a Toolbox for the Covalent Functionalization of Carbon Nanotubes with Peptides and Proteins. Angew. Chem. Int. Ed. 59, 17732–17738 (2020).

16. Cho, S.-Y. et al. Antibody-Free Rapid Detection of SARS-CoV-2 Proteins Using Corona Phase Molecular Recognition to Accelerate Development Time. Anal. Chem. 93, 14685–14693 (2021).

17. Landry, M. P. et al. Single-molecule detection of protein efflux from microorganisms using fluorescent single-walled carbon nanotube sensor arrays. Nat. Nanotechnol. 12, 368–377 (2017).

18. Pinals, R. L. et al. Rapid SARS-CoV-2 Spike Protein Detection by Carbon Nanotube-Based Near-Infrared Nanosensors. Nano Lett. 21, 2272–2280 (2021).

19. Bisker, G. et al. Protein-targeted corona phase molecular recognition. Nat. Commun. 7, 10241 (2016).

20. Pinals, R. L., Yang, D., Lui, A., Cao, W. & Landry, M. P. Corona Exchange Dynamics on Carbon Nanotubes by Multiplexed Fluorescence Monitoring. J. Am. Chem. Soc. 142, 1254– 1264 (2020).

21. Fu, F., Crespy, D., Landfester, K. & Jiang, S. In situ characterization techniques of protein corona around nanomaterials. Chem. Soc. Rev. 53, 10827–10851 (2024).

22. Shang, L. & Nienhaus, G. U. Fluorescent nanoparticle interactions with biological systems: What have we learned so far? in (eds Parak, W. J., Osinski, M. & Liang, X.-J.) 93380M (San Francisco, California, United States, 2015). doi:10.1117/12.2075722.

23. Shang, L. & Nienhaus, G. U. In Situ Characterization of Protein Adsorption onto Nanoparticles by Fluorescence Correlation Spectroscopy. Acc. Chem. Res. 50, 387–395 (2017).

24. Gao, L.-X. et al. Protein Labeling Facilitates the Understanding of Protein Corona Formation via Fluorescence Resonance Energy Transfer and Fluorescence Correlation Spectroscopy. Langmuir 39, 15275–15284 (2023).

25. Qu, S., Sun, F., Qiao, Z., Li, J. & Shang, L. In Situ Investigation on the Protein Corona Formation of Quantum Dots by Using Fluorescence Resonance Energy Transfer. Small 16, 1907633 (2020).

26. Tan, X. & Welsher, K. Particle‐by‐Particle In Situ Characterization of the Protein Corona via Real‐Time 3D Single‐Particle‐Tracking Spectroscopy**. Angew. Chem. Int. Ed. 60, 22359– 22367 (2021).

27. Antman-Passig, M. et al. Optical Nanosensor for Intracellular and Intracranial Detection of Amyloid-Beta. ACS Nano 16, 7269–7283 (2022).

28. Lee, S. et al. Clinically-Driven Rapidly Developed Nanoparticle Corona for Label-Free Cerebrospinal Fluid Leakage Detection. ACS Nano 19, 950–962 (2025).

29. Jin, X. et al. Corona Phase Molecular Recognition of the Interleukin-6 (IL-6) Family of Cytokines Using nIR Fluorescent Single-Walled Carbon Nanotubes. ACS Appl. Nano Mater. 6, 9791–9804 (2023).

30. Bisker, G. et al. Insulin Detection Using a Corona Phase Molecular Recognition Site on Single-Walled Carbon Nanotubes. ACS Sens. 3, 367–377 (2018).

31. Alizadehmojarad, A. A., Kang, H., Sánchez-Velázquez, G., Gong, X. & Strano, M. S. Quantum Defects Engineered in Chirality Pure Single-Wall Carbon Nanotubes for Enabling Ratiometric Sensing of Insulin. J. Am. Chem. Soc. 147, 40466–40480 (2025).

32. Bachilo, S. M. et al. Structure-Assigned Optical Spectra of Single-Walled Carbon Nanotubes. Science 298, 2361–2366 (2002).

33. Choi, J. H. & Strano, M. S. Solvatochromism in single-walled carbon nanotubes. Appl. Phys. Lett. 90, 223114 (2007).

34. O’Connell, M. J. et al. Band Gap Fluorescence from Individual Single-Walled Carbon Nanotubes. Science 297, 593–596 (2002).

35. Ouyang, M.Huang, J.-L. & Lieber, C. M. Fundamental Electronic Properties and Applications of Single-Walled Carbon Nanotubes. Acc. Chem. Res. 35, 1018–1025 (2002).

36. Porter, T. K., Sánchez-Velázquez, G. & Strano, M. S. The Role of Basal H_2_ O_2_ Concentration in ROS Stress Signaling Waveforms In Planta. ACS Agric. Sci. Technol. 5, 1434– 1441 (2025).

37. Ang, M. C.-Y. et al. Decoding early stress signaling waves in living plants using nanosensor multiplexing. Nat. Commun. 15, 2943 (2024).

38. Lew, T. T. S. et al. Real-time detection of wound-induced H2O2 signalling waves in plants with optical nanosensors. Nat. Plants 6, 404–415 (2020).

39. Jin, X. et al. A Nanosensor Platform for Biologging in Marine Animals. ACS Sens. 10, 4423–4433 (2025).

40. Kruss, S. et al. Neurotransmitter Detection Using Corona Phase Molecular Recognition on Fluorescent Single-Walled Carbon Nanotube Sensors. J. Am. Chem. Soc. 136, 713–724 (2014).

41. Strano, M. S. et al. Reversible, Band-Gap-Selective Protonation of Single-Walled Carbon Nanotubes in Solution. J. Phys. Chem. B 107, 6979–6985 (2003).

42. Cognet, L. et al. Stepwise Quenching of Exciton Fluorescence in Carbon Nanotubes by Single-Molecule Reactions. Science 316, 1465–1468 (2007).

43. Barone, P. W., Baik, S., Heller, D. A. & Strano, M. S. Near-infrared optical sensors based on single-walled carbon nanotubes. Nat. Mater. 4, 86–92 (2005).

44. Larsen, B. A. et al. Effect of Solvent Polarity and Electrophilicity on Quantum Yields and Solvatochromic Shifts of Single-Walled Carbon Nanotube Photoluminescence. J. Am. Chem. Soc. 134, 12485–12491 (2012).

45. Swenson, H. & Stadie, N. P. Langmuir’s Theory of Adsorption: A Centennial Review. Langmuir 35, 5409–5426 (2019).

46. Saptarshi, S. R., Duschl, A. & Lopata, A. L. Interaction of nanoparticles with proteins: relation to bio-reactivity of the nanoparticle. J. Nanobiotechnology 11, 26 (2013).

47. Gong, X. et al. Divalent Metal Cation Optical Sensing Using Single-Walled Carbon Nanotube Corona Phase Molecular Recognition. Anal. Chem. 94, 16393–16401 (2022).

48. Brege, J. J., Gallaway, C. & Barron, A. R. Fluorescence Quenching of Single-Walled Carbon Nanotubes with Transition-Metal Ions. J. Phys. Chem. C 113, 4270–4276 (2009).

49. Lu, Y., Yeung, N., Sieracki, N. & Marshall, N. M. Design of functional metalloproteins. Nature 460, 855–862 (2009).

50. Neuman, D. et al. Quantum Dot Fluorescence Quenching Pathways with Cr(III) Complexes. Photosensitized NO Production from trans -Cr(cyclam)(ONO)_2_^+^. J. Am. Chem. Soc. 130, 168–175 (2008).

51. Du Fossé, I., Boehme, S. C., Infante, I. & Houtepen, A. J. Dynamic Formation of Metal-Based Traps in Photoexcited Colloidal Quantum Dots and Their Relevance for Photoluminescence. Chem. Mater. 33, 3349–3358 (2021).

52. Masuhara, H. et al. Fluorescence quenching mechanism of aromatic hydrocarbons by closed-shell heavy metal ions in aqueous and organic solutions. J. Phys. Chem. 88, 5868–5873 (1984).

53. Kim, I. J., Xu, Y. & Nam, K. H. Metal-Induced Fluorescence Quenching of Photoconvertible Fluorescent Protein DendFP. Molecules 27, 2922 (2022).

54. Chakraborti, S. & Chakrabarti, P. Self-Assembly of Ferritin: Structure, Biological Function and Potential Applications in Nanotechnology. in Biological and Bio-inspired Nanomaterials (eds Perrett, S., Buell, A. K. & Knowles, T. P. J.) vol. 1174 313–329 (Springer Singapore, Singapore, 2019).

55. Erickson, H. P. Size and Shape of Protein Molecules at the Nanometer Level Determined by Sedimentation, Gel Filtration, and Electron Microscopy. Biol. Proced. Online 11, 32–51 (2009).

56. Ma, X. et al. Single-Walled Carbon Nanotubes Alter Cytochrome c Electron Transfer and Modulate Mitochondrial Function. ACS Nano 6, 10486–10496 (2012).

57. Yang, M. et al. Carbon Nanotubes Induce Secondary Structure Changes of Bovine Albumin in Aqueous Phase. J. Nanosci. Nanotechnol. 10, 7550–7553 (2010).

58. Lou, K. et al. Comprehensive studies on the nature of interaction between carboxylated multi-walled carbon nanotubes and bovine serum albumin. Chem. Biol. Interact. 243, 54–61 (2016).

59. Wang, Y.-Q.Zhang, H.-M. & Cao, J. Binding of hydroxylated single-walled carbon nanotubes to two hemoproteins, hemoglobin and myoglobin. J. Photochem. Photobiol. B 141, 26–35 (2014).

60. Nelson, J. T. et al. Mechanism of Immobilized Protein A Binding to Immunoglobulin G on Nanosensor Array Surfaces. Anal. Chem. 87, 8186–8193 (2015).

61. Ledesma, F. et al. Covalent Attachment of Horseradish Peroxidase to Single‐Walled Carbon Nanotubes for Hydrogen Peroxide Detection. Adv. Funct. Mater. 34, 2316028 (2024).

62. Ignatenko, O. V., Rubtsova, M. Y., Ivanova, N. L., Ouporov, I. V. & Egorov, A. M. ANALYSIS OF THE ANTIGENIC STRUCTURE OF HOLO-AND APO-FORMS OF HORSERADISH PEROXIDASE.

63. Cha, H. J., Jang, D. S., Jin, K. S. & Choi, K. Y. Structural analyses combined with small-angle X-ray scattering reveals that the retention of heme is critical for maintaining the structure of horseradish peroxidase under denaturing conditions. Amino Acids 49, 715–723 (2017).

64. Silmore, K. S., Gong, X., Strano, M. S. & Swan, J. W. High-Resolution Nanoparticle Sizing with Maximum A Posteriori Nanoparticle Tracking Analysis. ACS Nano 13, 3940–3952 (2019).

65. Sánchez-Velázquez, G. et al. Using Molecular Probe Adsorption to Characterize the Nanoparticle Corona Phase and Molecular Recognition. Langmuir 41, 17602–17614 (2025).

66. Park, M. et al. Measuring the Accessible Surface Area within the Nanoparticle Corona Using Molecular Probe Adsorption. Nano Lett. 19, 7712–7724 (2019).

