## Supporting Information for "Solution Phase Protein Adsorption to ss(GT)_15_-DNA Wrapped Single Walled Carbon Nanotubes"

### Calibration Curves for all Proteins:

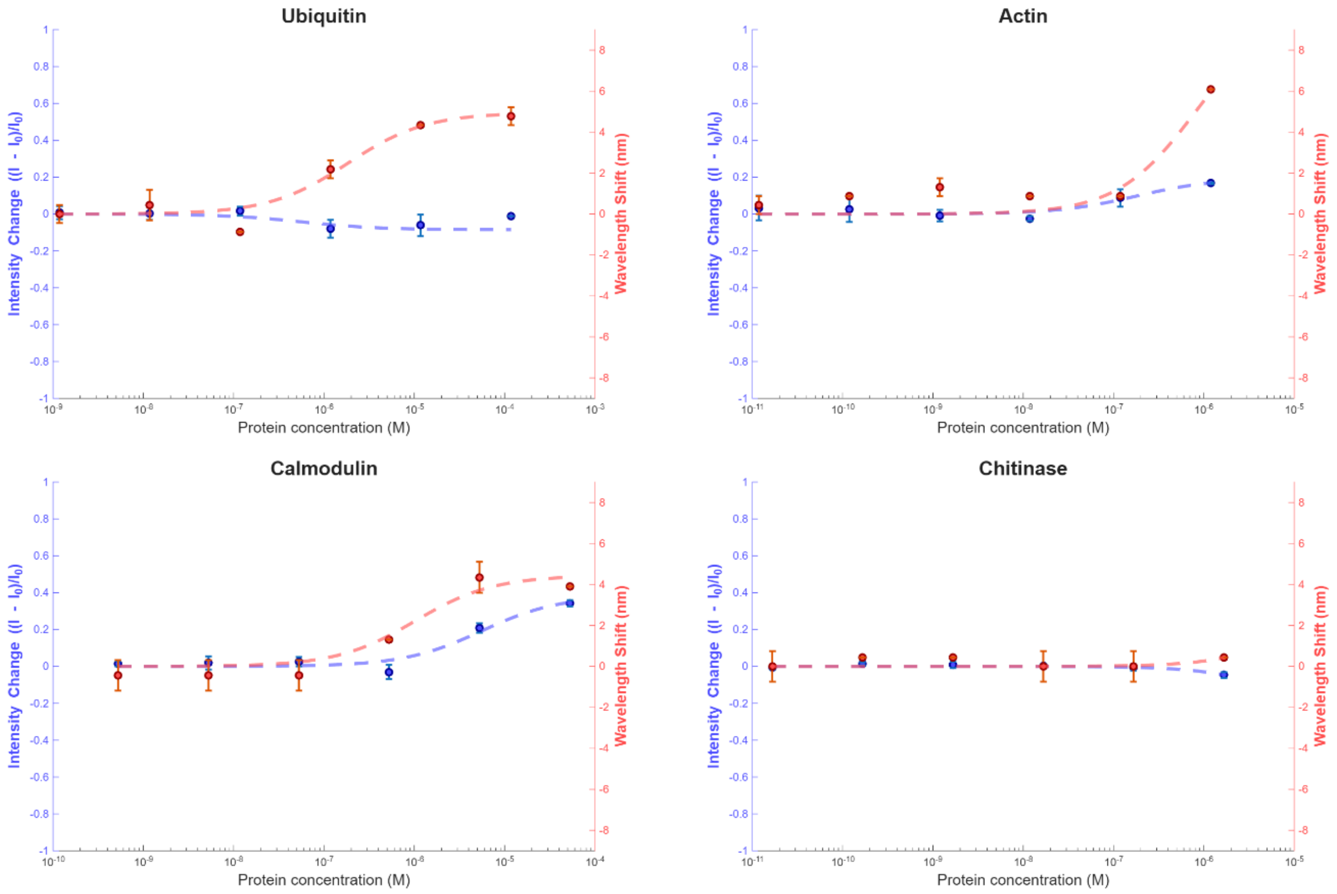

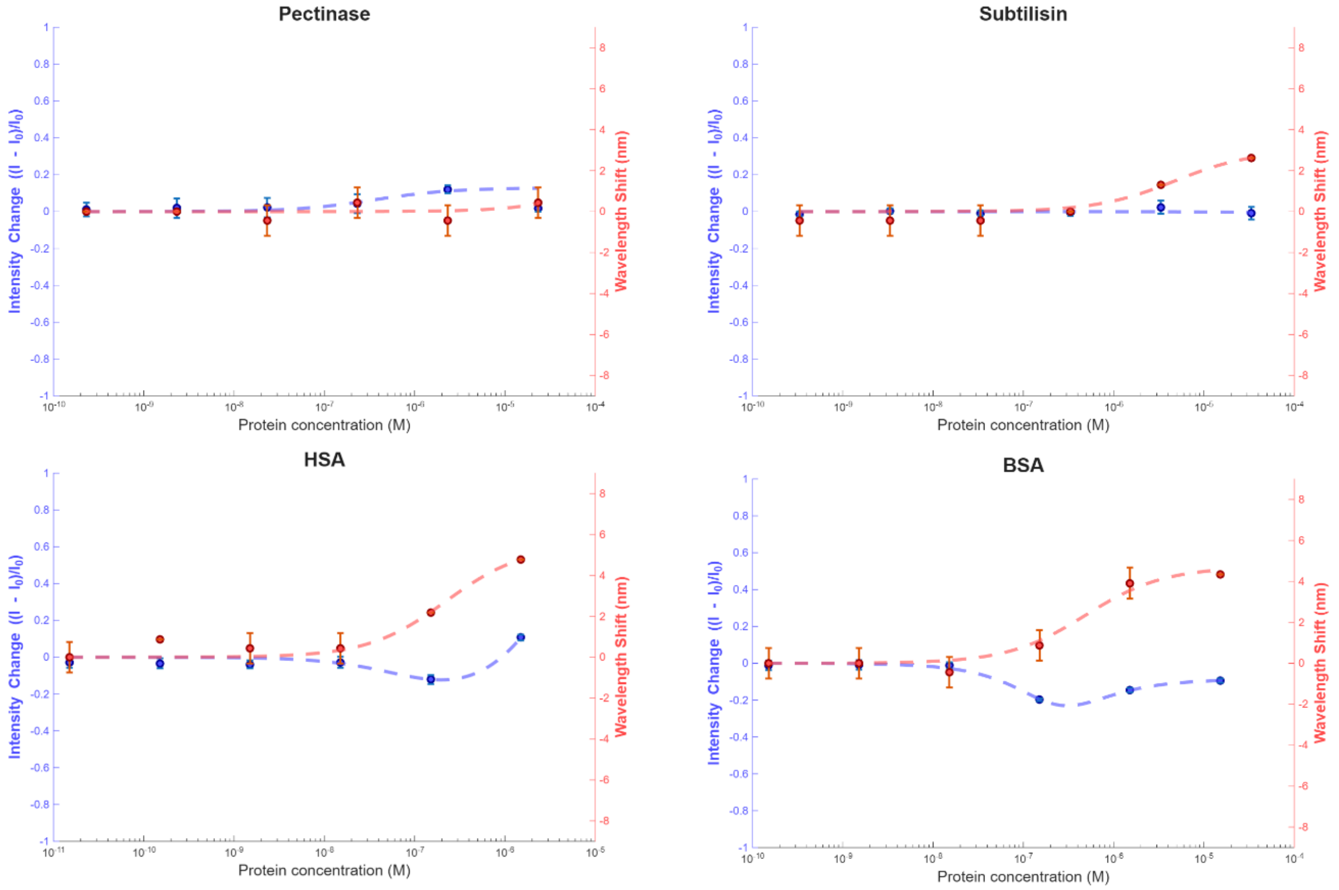


**Supporting Figure 1**. Calibration curves for non-metal–containing proteins. Dashed lines indicate fits to either the single-site Langmuir isotherm or the two-step adsorption model, as appropriate. The left y-axis shows the normalized intensity change, and the right y-axis shows the corresponding wavelength shift.

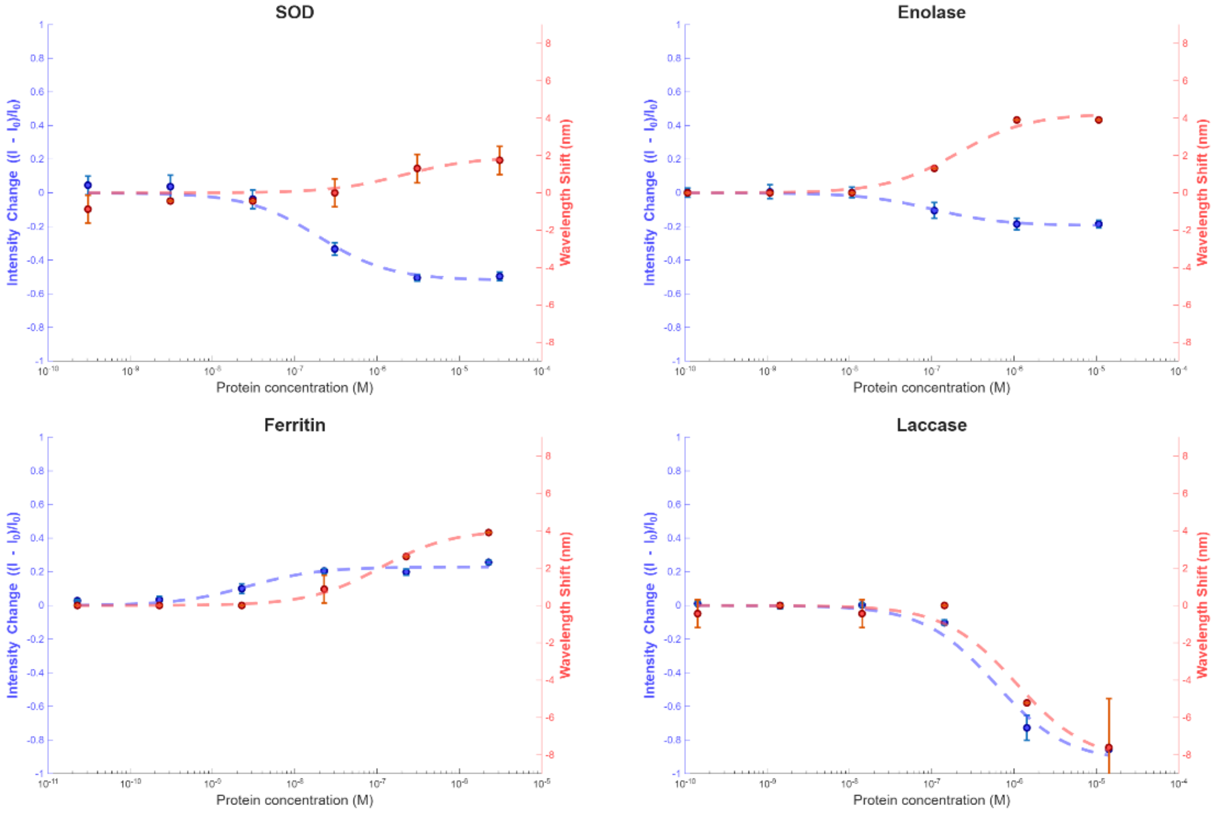

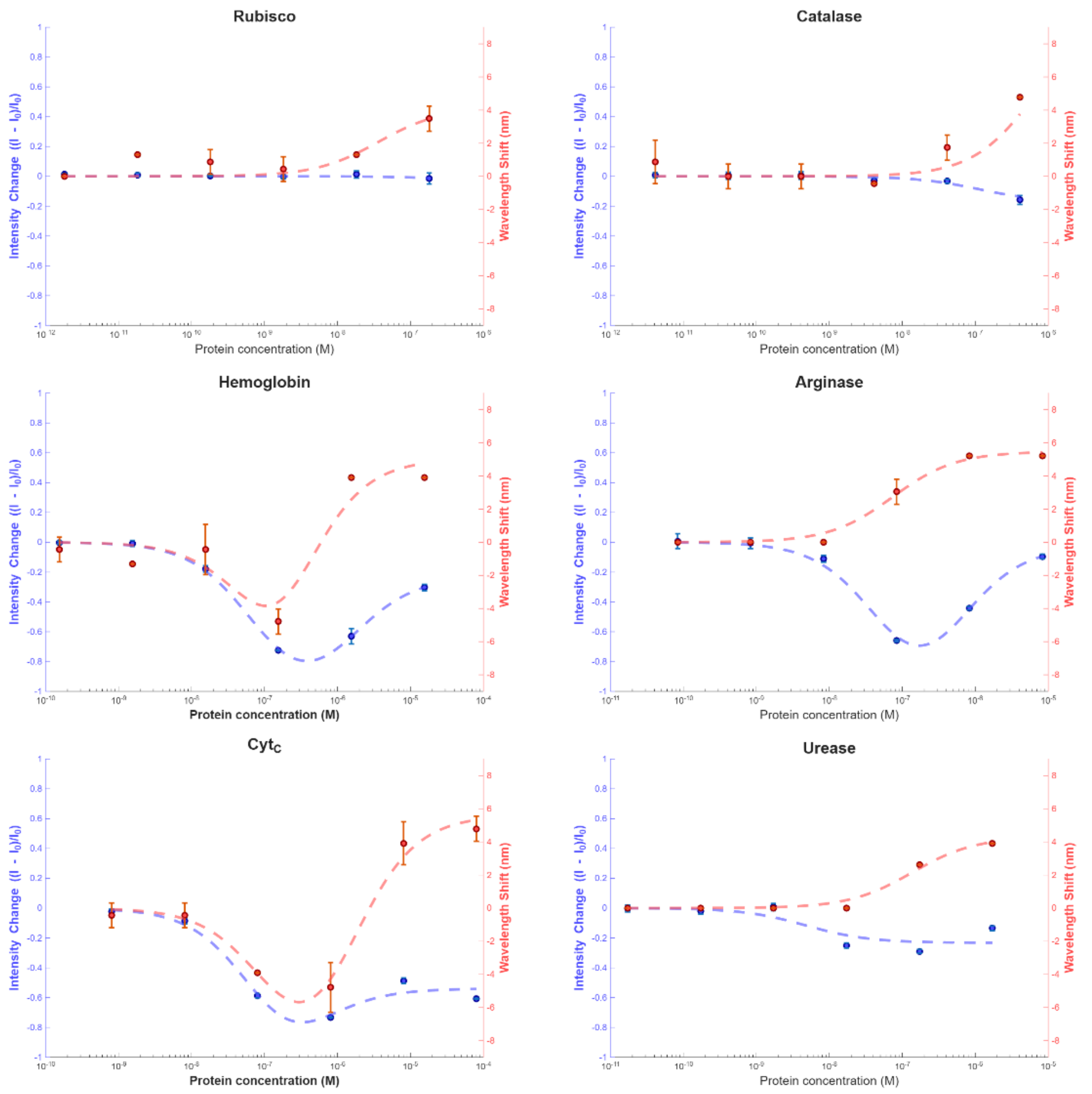


**Supporting Figure 2**. Calibration curves for metal–containing proteins. Dashed lines indicate fits to either the single-site Langmuir isotherm or the two-step adsorption model, as appropriate. The left y-axis shows the normalized intensity change, and the right y-axis shows the corresponding wavelength shift.

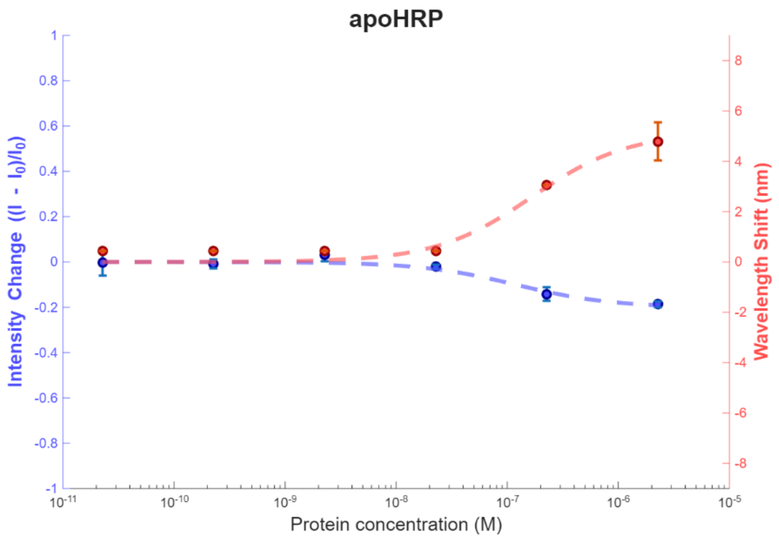


**Supporting Figure 3**. Calibration curves for apo-HRP. Dashed lines indicate fits to the single-site Langmuir isotherm model. The left y-axis shows the normalized intensity change, and the right y-axis shows the corresponding wavelength shift.

### Fitted Parameters for Protein Calibration Curves:

**Supporting Table 1:** Fitted Parameters and $R^2$ for single-site Langmuir isotherm model for the intensity channel.

| Protein | $K_I$ (mg/mL) | $R^2$ | β |
|---|---|---|---|
| ADH | 0.048 | 0.9698 | 0.3868 |
| Actin | 0.0075 | 0.8799 | 0.1941 |
| Calmodulin | 0.1057 | 0.9518 | 0.3847 |
| SOD | 0.0063 | 0.9842 | -0.5196 |
| Enolase | 0.0096 | 0.9851 | -0.1948 |
| Ferritin | 0.0012 | 0.9586 | 0.229 |
| HRP | 4.05E-05 | 0.9752 | -0.9561 |
| Catalase | 0.061 | 0.9643 | -0.2544 |
| Laccase | 0.0412 | 0.9848 | -0.9278 |
| Apo-HRP | 0.0053 | 0.9639 | -0.2011 |
| Urease | 0.0028 | 0.7625 | -0.2326 |

**Supporting Table 2:** Fitted Parameters and $R^2$ for single-site Langmuir isotherm model for the wavelength channel.

| Protein | $K_I$ (mg/mL) | $R^2$ | β |
|---|---|---|---|
| Ubiquitin | 0.0156 | 0.9478 | 4.9474 |
| ADH | 0.0936 | 0.9716 | 5.6156 |
| Actin | 0.034 | 0.8706 | 10 |

| | | | |
|---|---|---|---|
| Subtilisin | 0.143 | 0.9221 | 2.9996 |
| Calmodulin | 0.0194 | 0.9454 | 4.4365 |
| BSA | 0.0323 | 0.9769 | 4.702 |
| HSA | 0.0148 | 0.9488 | 5.4827 |
| SOD | 0.0679 | 0.7771 | 1.9223 |
| Enolase | 0.0191 | 0.9876 | 4.2376 |
| Ferritin | 0.0485 | 0.9964 | 4.0492 |
| Urease | 0.0797 | 0.9823 | 4.3032 |
| Arginase | 0.0089 | 0.9891 | 5.4721 |
| HRP | 7.79E-05 | 0.9492 | -8.061 |
| Rubisco | 0.0201 | 0.7131 | 4.1673 |
| Catalase | 0.028 | 0.9403 | 6.1388 |
| Laccase | 0.0746 | 0.9747 | -8.3434 |
| apo-HRP | 0.0073 | 0.9726 | 5.1571 |

**Supporting Table 3:** Fitted Parameters and $R^2$ for two-step adsorption model for the intensity channel.

| Protein | $K_{1,I}$ (mg/mL)$^{-1}$ | $K_{2,I}$ (mg/mL)$^{-1}$ | $\beta_1$ | $\beta_2$ | $R^2$ |
|---|---|---|---|---|---|
| BSA | 60.95988 | 60.84735 | -0.57349 | -0.08422 | 0.987892 |
| HSA | 168.9829 | 3.504625 | -0.23456 | 1.099984 | 0.885806 |
| Hemoglobin | 213.0955 | 10.86917 | -1.1 | -0.23005 | 0.998927 |
| CTC | 1125.408 | 181.89 | -1.09998 | -0.53925 | 0.979318 |
| Arginase | 165.9245 | 14.8507 | -1.1 | -0.0269 | 0.992393 |

**Supporting Table 4:** Fitted Parameters and $R^2$ for two-step adsorption model for the wavelength channel.

| Protein | $K_{1,\lambda}$ (mg/mL)$^{-1}$ | $K_{2,\lambda}$ (mg/mL)$^{-1}$ | $\beta_1$ | $\beta_2$ | $R^2$ |
|---|---|---|---|---|---|
| Hemoglobin | 185.1197 | 51.51909 | -10 | 5.042471 | 0.883035 |
| CTC | 711.1564 | 51.8972 | -10 | 5.620252 | 0.981422 |

### Parity plot between dissociation constants derived from intensity ($K_{D,I}$) and wavelength ($K_{D,\lambda}$):

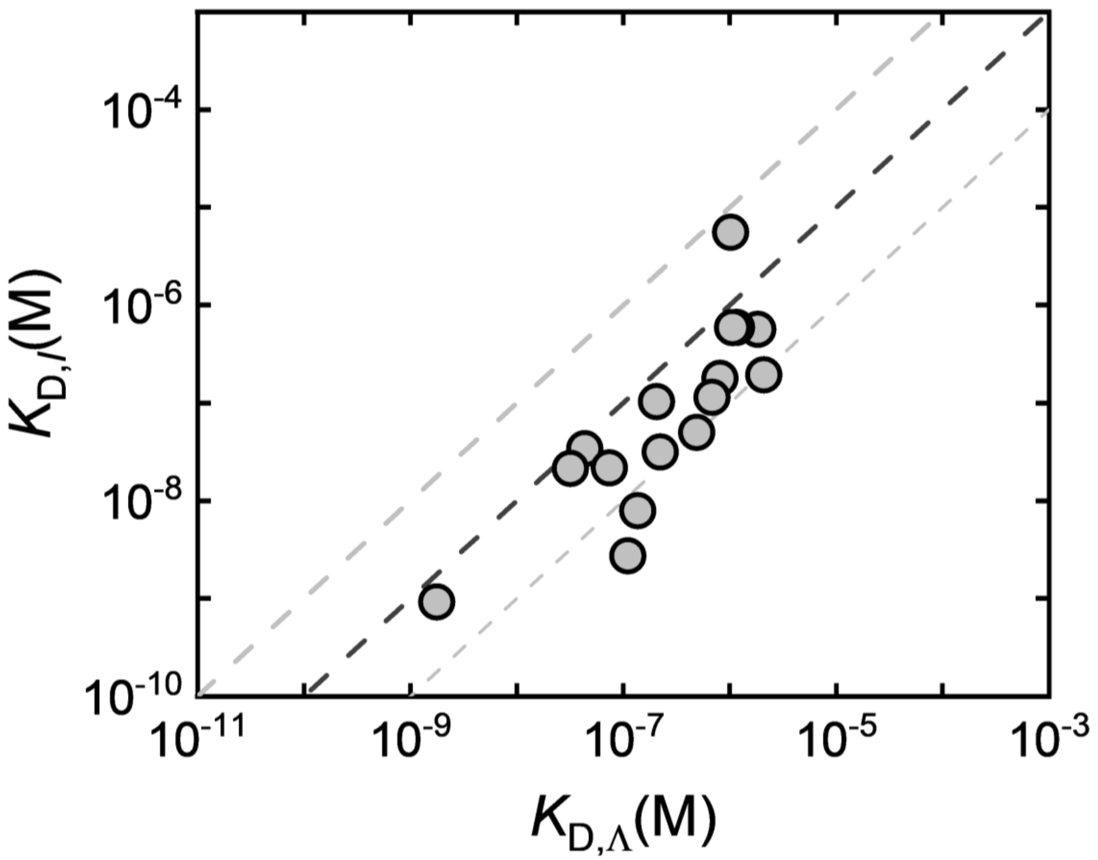


**Supporting Figure 4**. Correlation between dissociation constants derived from intensity ($K_{D,I}$) and wavelength ($K_{D,\lambda}$). Dashed grey lines mark ±1 $\log_{10}$ deviation from the 1:1 parity line (dashed black line).

### Equilibrium Equivalence of Surface-Association and Solution-Association Two-Step Models

In addition to the sequential surface-binding scheme described in the main text, an equivalent equilibrium dependence is obtained if proteins first associate in solution and the resulting complex subsequently adsorbs to the nanotube surface. In this alternative mechanism, proteins reversibly form a higher-order complex in bulk solution, followed by adsorption of that complex to an available surface site:

$$P + P \overset{K_P}{\leftrightarrow} PP \quad (S1)$$

$$\theta + PP \overset{K_3}{\leftrightarrow} \theta_{PP} \quad (S2)$$

$$\theta_{PP} = K_3 K_P \theta [P]^2 \quad (S3)$$

Comparing this expression to the two-step surface scheme in the main text, where $\theta_{PP} = K_1 K_2 \theta [P]^2$, shows that the equilibrium behavior is identical provided that

$$K_3 K_P = K_1 K_2 = K_{eff} \quad (S4)$$

Here, $K_{\text{eff}}$ is an effective second-order equilibrium constant that captures the net propensity to form the $\theta_{PP}$ state, whether that state arises from sequential adsorption at the surface (protein binding to a pre-occupied site) or from solution-phase protein association followed by adsorption. Accordingly, at equilibrium, the two-step model cannot uniquely distinguish between these microscopic pathways, and $K_1K_2$ is interpreted as an effective parameter encompassing both mechanisms.

**Inter-User and Batch-to-Batch Reproducibility of SWCNT–holo-HRP Binding:**

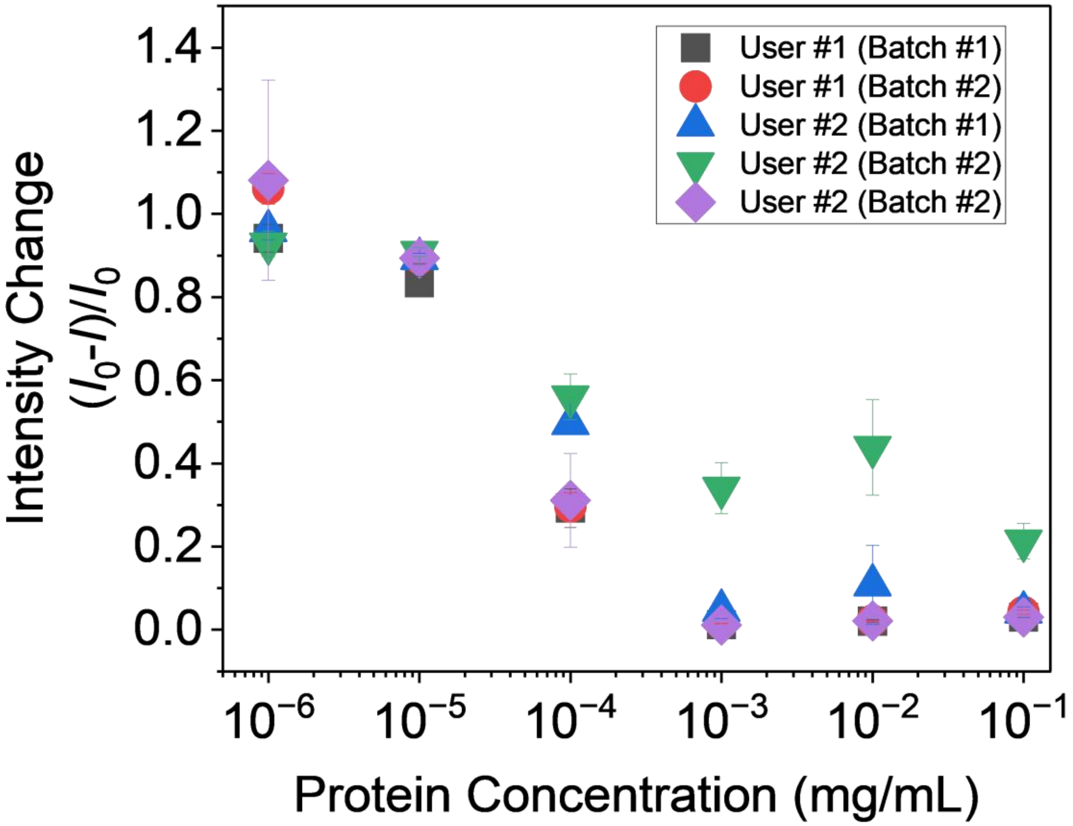


**Supporting Figure 5. Batch-to-batch reproducibility of holo-HRP titrations on SWCNTs.** Fluorescence quenching response as a function of holo-HRP concentration for independently prepared SWCNT batches produced by two users (User #1 and User #2; batches prepared on different days). Despite minor differences in quenching magnitude between batches, fitted affinities were similar across preparations: User #1 (Batch #1) $K_d = 1.13 \times 10^{-9}$ M; User #1 (Batch #2) $K_d = 1.10 \times 10^{-9}$ M; User #2 (Batch #1) $K_d = 1.83 \times 10^{-9}$ M; User #2 (Batch #2) $K_d = 1.36 \times 10^{-9}$ M; and User #2 (Batch #2, repeat) $K_d = 1.30 \times 10^{-9}$ M.

**Molecular Probe Adsorption:**

Molecular probe adsorption (MPA) quantifies the solvent-accessible nanoparticle surface area by leveraging a fluorescent probe (riboflavin) that is quenched upon adsorption to exposed SWCNT surface sites. A known SWCNT concentration ($C_{NP}$) is titrated with increasing total probe concentrations ($C_{total}$); after equilibration, the concentration of unbound probe remaining in solution ($C_{probe}$) is determined from measured fluorescence using a probe calibration curve. Assuming Langmuir adsorption, the probe material balance yields a linear form:

$$C_{NP}/\Delta = 1/q + (K_D'/q)(1/C_{probe}),\ \Delta = C_{total} - C_{probe}$$

where $q$ is the maximum amount of probe that can be adsorbed per unit mass (or mole) of nanoparticle, and $K_D'$ is the probe–SWCNT dissociation constant. Thus, a plot of $C_{NP}/\Delta$ versus $1/C_{probe}$ is linear with slope $K_D'/q$, and the reciprocal parameter $q/K_D'$ serves as a metric of accessible SWCNT surface area (smaller $q/K_D$indicates lower accessibility).

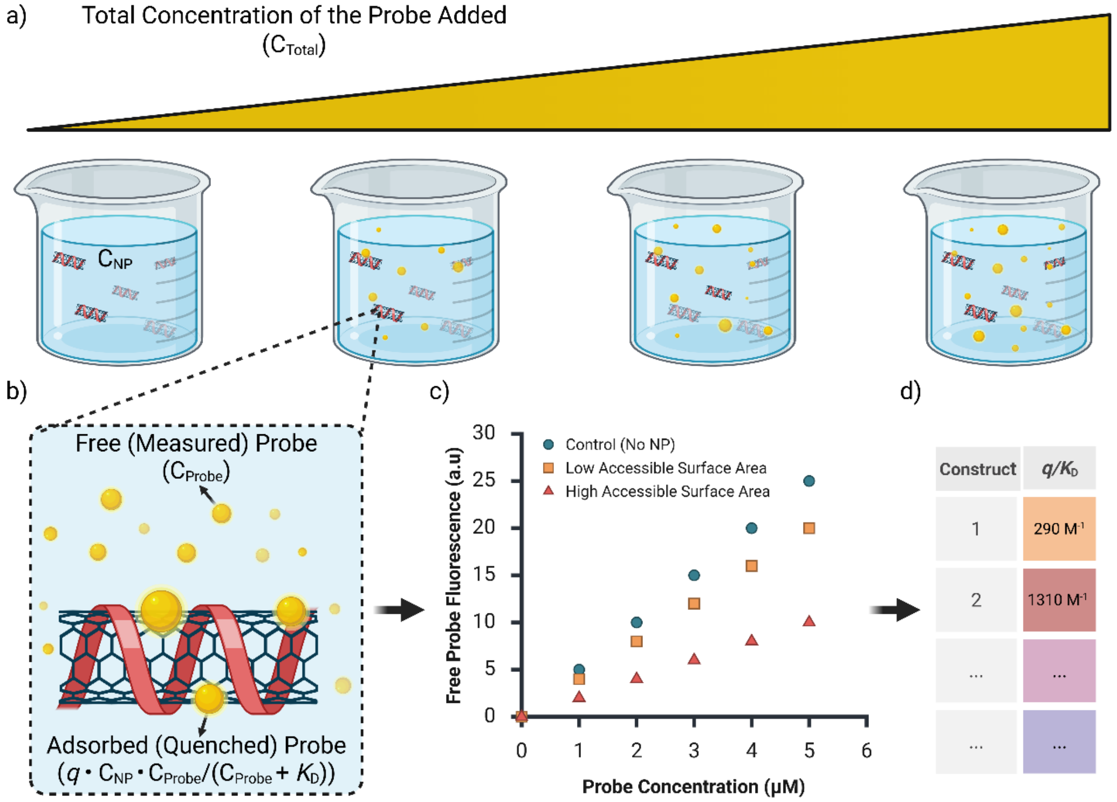


**Supporting Figure 6.** Schematic representation of the molecular probe adsorption (MPA) technique. (a) Illustration of a titration experiment in which a fluorescent dye (probe) is added across samples containing the same underlying nanoparticle concentration $C_{NP}$. (b) In each sample, the probe exists in solution at a measurable concentration $C_{probe}$, while a portion adsorbs to exposed (i.e., non–corona-shielded) regions of the nanoparticle surface, where it is quenched. The parameter $q$ denotes the number of probe molecules adsorbed per unit mass of nanoparticle, and $K_D'$ is the dissociation constant describing the probe–nanoparticle interaction. (c) Free probe fluorescence is measured using a microplate reader. Nanoparticle constructs with low accessible surface area (orange) result in higher free probe fluorescence, while constructs with high accessible surface area (red) show greater probe quenching and thus lower fluorescence. (d) The MPA analysis enables calculation of the $q/K_D'$ parameter for different nanoparticle–corona constructs based on the measured free probe fluorescence, the total probe added, and the known nanoparticle concentration. This allows users to quantitatively compare constructs: lower $q/K_D'$ values correspond to lower accessible surface area, while higher values indicate greater probe accessibility to the nanoparticle surface. Created in BioRender. Sánchez-Velázquez, G. (2025) https://BioRender.com/y2x65a8. Reproduced with permission from Sánchez-Velázquez, Gabriel, et al. "Using Molecular Probe Adsorption to Characterize the Nanoparticle Corona Phase and Molecular Recognition." *Langmuir* (2025).

In practice, MPA requires two fluorescence calibration curves acquired on a plate reader: (i) probe in buffer (to convert fluorescence to $C_{probe}$) and (ii) probe in the presence of the nanoparticle construct to quantify the unbound probe after quenching by accessible SWCNT surface sites. In this study, MPA was performed on G-SWCNT dispersions incubated with apo-HRP or holo-HRP (or no protein) to assess whether protein adsorption altered accessible SWCNT surface area. Because free ssDNA was not removed prior to these measurements, and because soluble HRP may also interact with riboflavin, probe quenching can include a background contribution not attributable to SWCNT adsorption. To account for this, matched background controls (probe with the relevant soluble components—free ssDNA and, where applicable, protein—but without SWCNT) were measured and incorporated via a background correction term ($C_{background}$).

Accordingly, the probe balance is written as:

$$C_{total} = C_{probe} + q_{background}\, C_{background} \frac{C_{probe}}{C_{probe}+K_{D,background}} + q_{NP}\, C_{NP} \frac{C_{probe}}{C_{probe}+K_D'}$$

$$\Delta = C_{total} - C_{probe} - q_{background}\, C_{background} \frac{C_{probe}}{K_{D,background}} = q_{NP}\, C_{NP} \frac{C_{probe}}{C_{probe}+K_D'}$$

$$\frac{C_{NP}}{\Delta} = \frac{1}{q_{NP}} + \frac{K_D'}{q_{NP}} \frac{1}{C_{probe}}$$

The background terms ($q_{background}$, $K_{D,background}$, and $C_{background}$) account for probe interactions with soluble species present in the assay (e.g., free ssDNA and, where applicable, HRP) that can contribute to quenching independent of SWCNT surface adsorption. Supplementary Figure 7 shows the fluorescence calibration data (buffer control, background, and G-SWCNT ± protein) used to determine $\frac{q}{K_D'}$.

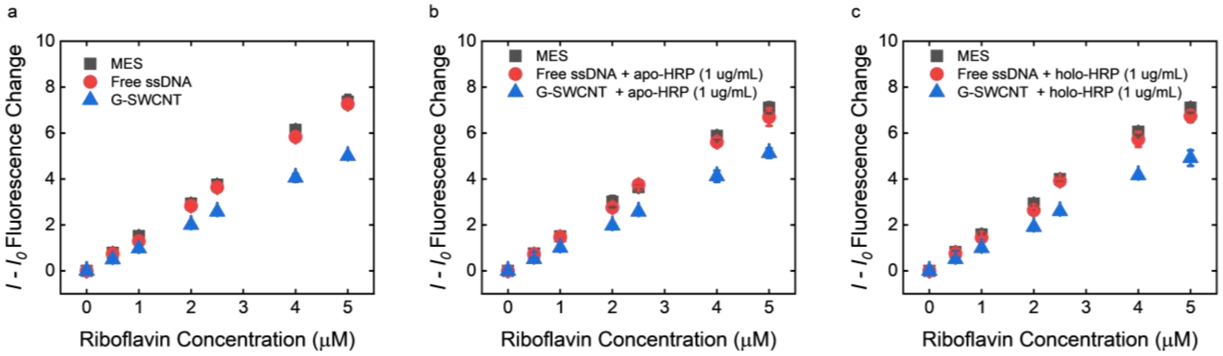


**Supporting Figure 7.** Probe fluorescence as a function of concentration in three experimental conditions: (a) no protein added, (b) apo-HRP (final concentration 1 μg/mL), and (c) holo-HRP (final concentration 1 μg/mL). Each experimental run includes a control (buffer only), a background condition (free ssDNA with or without protein, as applicable), and the G-SWCNT construct with protein added where indicated.

### Zeta Potential:

G-SWCNT zeta potential was -30.47 ± 2.00 mV, measured at 5 mg/L in 10 mM KCl + 10 mM Tris-EDTA buffer at pH 8.0. The reported value is average ± standard deviation from three measurements with 10 cycles each. Detailed measurement conditions and model fitting information for each measurement are below (**Supporting Table 5**, **Supporting Figures 8-10**).

**Supporting Table 5.** Zeta Potential Replicates

| Zeta Potential (mV) | Mobility (μ/s)/(V/cm) | Conductance (μS) | Sample Count Rate (kcps) | Ref. Count Rate (kcps) | RMS Residual |
|---|---|---|---|---|---|
| -31.92 | -2.49 | 5,201 | 301 | 632 | 1.7931E-02 |
| -28.18 | -2.20 | 5,201 | 301 | 632 | 4.8626E-02 |
| -31.30 | -2.45 | 5,208 | 577 | 729 | 5.4163E-02 |

Brookhaven Instruments

Advanced PALS Report

| Sample | | Results | |
|---|---|---|---|
| Type: | PALS | Zeta Potential (mV): | -31.30 |
| Sample ID: | XW_20260216_GT15_TKP_10mMKCl_TE8 - 4 | Zeta Potential Model: | Smoluchowski |
| Operator ID: | Unknown Operator | Mobility (μ/s)/(V/cm): | -2.45 |
| SOP ID: | Xiaoyi | Cycles Collected: | 10 |
| Start Date/Time: | 2/16/2026 7:08:30 PM | **Measurement Conditions** | |
| **Advanced ID** | | Conductance (μS): | 5,208 |
| Project ID: | | Voltage Applied (V): | 4.00 |
| Group ID: | | Wavelength (nm): | 640.0 |
| Batch #: | 0 | Electric Field (V/cm): | 9.81 |
| Rev #: | 0 | Field Frequency (Hz): | 2.00 |
| Revision Date/Time: | | RMS Residual: | 5.4163e-02 |
| **Liquid** | | Ref. Count Rate (kcps): | 729 |
| Liquid: | Water | Sample Count Rate (kcps): | 577 |
| Temp. (°C): | 25.00 | Count Rate Ratio: | 1.26 |
| Viscosity (cP): | 0.8900 | Simulated Data: | ☐ |
| Ref. Index: | 1.3310 | **Cell & Electrode** | |
| Dielectric Constant: | 78.54 | Cell Description: | BI-SCP |
| pH: | 8.00 | Electrode Assembly: | BI-ZEL (1,250 μL) |

Notes:

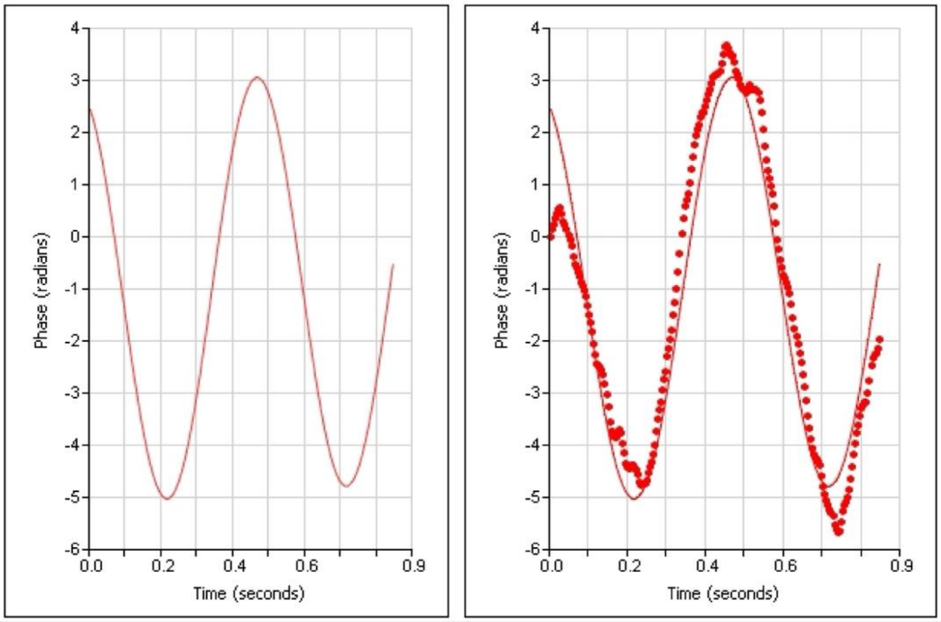


Printed: 2/19/2026 Particle Solutions v. 3.5 (Massachusetts Institute of Technology) 1 of 1

**Supporting Figure 8.** Details of zeta potential measurement for first measurement.

Brookhaven Instruments

### Advanced PALS Report

| Sample | | Results | |
|---|---|---|---|
| Type: | PALS | Zeta Potential (mV): | -31.92 |
| Sample ID: | XW_20260216_GT15_TKP_10mMKCl_TE8 - 1 | Zeta Potential Model: | Smoluchowski |
| Operator ID: | Unknown Operator | Mobility (μ/s)/(V/cm): | -2.49 |
| SOP ID: | Xiaoyi | Cycles Collected: | 10 |

| Sample | |
|---|---|
| Start Date/Time: | 2/16/2026 6:55:11 PM |

| Advanced ID | |
|---|---|
| Project ID: | |
| Group ID: | |
| Batch #: | 0 |
| Rev #: | 0 |
| Revision Date/Time: | |

| Liquid | |
|---|---|
| Liquid: | Water |
| Temp. (°C): | 25.00 |
| Viscosity (cP): | 0.8900 |
| Ref. Index: | 1.3310 |
| Dielectric Constant: | 78.54 |
| pH: | 8.00 |

| Measurement Conditions | |
|---|---|
| Conductance (μS): | 5,201 |
| Voltage Applied (V): | 4.00 |
| Wavelength (nm): | 640.0 |
| Electric Field (V/cm): | 9.81 |
| Field Frequency (Hz): | 2.00 |
| RMS Residual: | 1.7931e-02 |
| Ref. Count Rate (kcps): | 632 |
| Sample Count Rate (kcps): | 301 |
| Count Rate Ratio: | 2.10 |
| Simulated Data: | ☐ |

| Cell & Electrode | |
|---|---|
| Cell Description: | BI-SCP |
| Electrode Assembly: | BI-ZEL (1,250 μL) |

Notes:

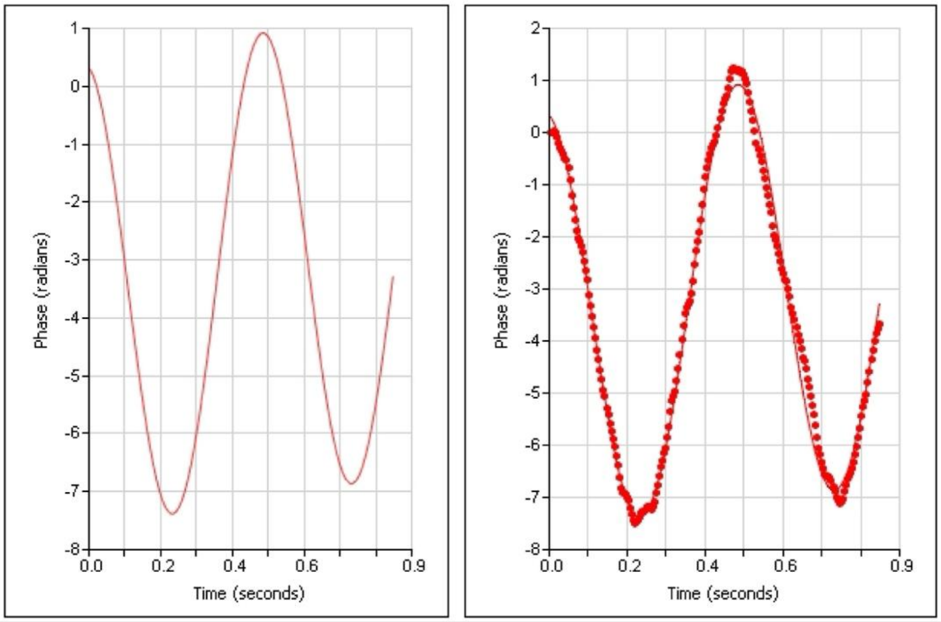


Printed: 2/19/2026 Particle Solutions v. 3.5 (Massachusetts Institute of Technology) 1 of 1

**Supporting Figure 9.** Details of zeta potential measurement for second measurement.

Brookhaven Instruments

Advanced PALS Report

| Sample | | Results | |
|---|---|---|---|
| Type: | PALS | Zeta Potential (mV): | -28.18 |
| Sample ID: | XW_20260216_GT15_TKP_10mMKCl_TE8 - 2 | Zeta Potential Model: | Smoluchowski |
| Operator ID: | Unknown Operator | Mobility (μ/s)/(V/cm): | -2.20 |
| SOP ID: | Xiaoyi | Cycles Collected: | 10 |
| Start Date/Time: | 2/16/2026 6:56:49 PM | **Measurement Conditions** | |
| **Advanced ID** | | Conductance (μS): | 5,201 |
| Project ID: | | Voltage Applied (V): | 4.00 |
| Group ID: | | Wavelength (nm): | 640.0 |
| Batch #: | 0 | Electric Field (V/cm): | 9.81 |
| Rev #: | 0 | Field Frequency (Hz): | 2.00 |
| Revision Date/Time: | | RMS Residual: | 4.8626e-02 |
| **Liquid** | | Ref. Count Rate (kcps): | 632 |
| Liquid: | Water | Sample Count Rate (kcps): | 301 |
| Temp. (°C): | 25.00 | Count Rate Ratio: | 2.10 |
| Viscosity (cP): | 0.8900 | Simulated Data: | ☐ |
| Ref. Index: | 1.3310 | **Cell & Electrode** | |
| Dielectric Constant: | 78.54 | Cell Description: | BI-SCP |
| pH: | 8.00 | Electrode Assembly: | BI-ZEL (1,250 μL) |

Notes:

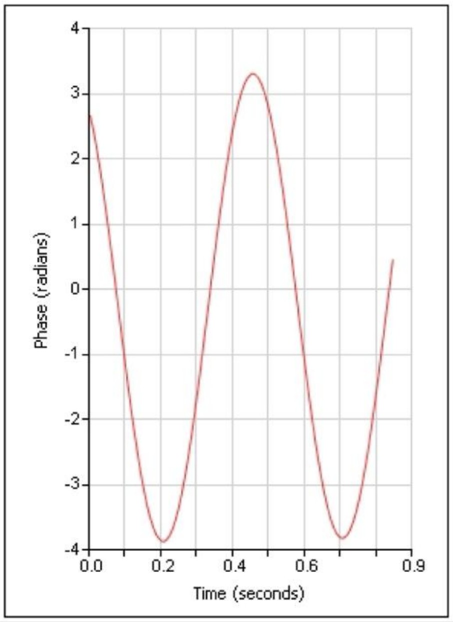


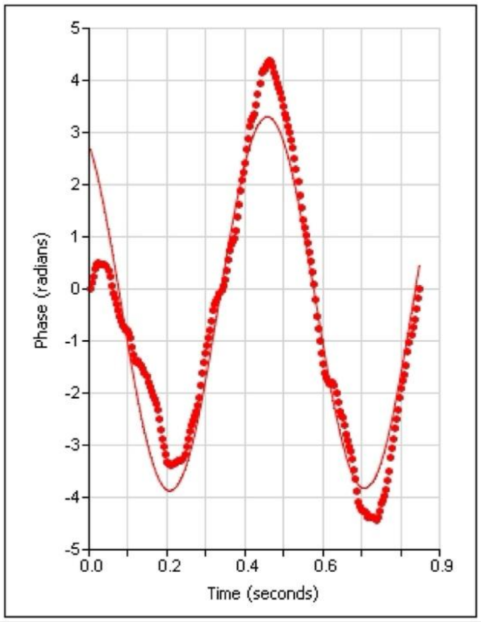




**Supporting Figure 10.** Details of zeta potential measurement for third measurement.
